# Salicylic acid-driven association of LENRV and NIMIN1/NIMIN2 binding domain regions in the C-terminus of tobacco NPR1 transduces SAR signal

**DOI:** 10.1101/543645

**Authors:** David Neeley, Evelyn Konopka, Anna Straub, Felix Maier, Artur J. P. Pfitzner, Ursula M. Pfitzner

## Abstract

- NONEXPRESSOR OF PATHOGENESIS-RELATED (PR) GENES1 (NPR1) is the central regulator of salicylic acid (SA)-induced *PR-1* gene expression and systemic acquired resistance (SAR). The mechanism how SA is transduced through NPR1 is discussed controversially. Previously, we showed that Arabidopsis and tobacco (Nt) NPR1 contain two domains in their C-terminal thirds with relevance to SA signaling. SA sensitivity of NPR1 relies on the arginine residue in the LENRV motif, and SA-induced NIM1-INTERACTING (NIMIN, N) proteins bind to a highly conserved sequence termed N1/N2 binding domain (BD).
- We demonstrate that LENRV and N1/N2BD regions of tobacco NPR1, separated from each other, interact in yeast, in vitro, in plant and in animal cells. Physical association of LENRV and N1/N2BD parts is enhanced considerably by SA and functional analogs, but not by a non-functional analog. Furthermore, physical association requires R431 and is most effective with intact LENRV and N1/N2BD interfaces.
- Association of separated LENRV and N1/N2BD parts by SA reconstitutes a functional NtNPR1 C-terminus, displaying transcription activity and able to interact with TGA transcription factors at two distinct sites.
- Tobacco NIMIN proteins can assemble LENRV and N1/N2BD parts into ternary complexes suggesting that NIMINs shape the NPR1 C-terminus to modulate SA signaling.

## Introduction

Plants have evolved different layers of defense to recognize and combat invading microbes (Jones & Dangl, 2006). The immune response systemic acquired resistance (SAR) becomes effective in non-infected leaves away from pathogen invasion without displaying macroscopic symptoms (Ross, 1961; Fu & Dong, 2013). The signal to induce SAR is salicylic acid (SA; Malamy *et al.*, 1990; Métraux *et al.*, 1990; Vlot *et al.*, 2009). In non-infected leaves, increasing levels of SA are paralleled by accumulation of PATHOGENESIS-RELATED (PR) proteins (Ward *et al.*, 1991; van Loon *et al.*, 2006), and exogenous application of SA or functional analogs is sufficient to elicit the resistance response and to induce *PR-1* genes that serve as markers for SAR (White, 1979; Ward *et al.*, 1991).

The central regulator of SAR is *NONEXPRESSOR OF PR GENES1* (*NPR1*), also known as *NON-INDUCIBLE IMMUNITY1* (*NIM1*) and *SALICYLIC ACID INSENSITIVE1* (*SAI1*) (Cao *et al.*, 1997; Ryals *et al.*, 1997; Shah *et al.*, 1997). Overexpression experiments strongly suggested that NPR1 is active only after SA challenge (Cao *et al.*, 1998; Friedrich *et al.*, 2001). NPR1 interacts with two groups of proteins. TGACG-BINDING FACTORS (TGAs) link NPR1 with SA responsive *as-1*-like *cis*-acting elements present in promoters of *PR-1* genes from tobacco (Nt) and Arabidopsis (At; Lebel *et al.*, 1998; Strompen *et al.*, 1998; Zhang *et al.*, 1999; Després *et al.*, 2000; Niggeweg *et al.*, 2000; Zhou *et al.*, 2000). Consistently, NPR1 proteins from Arabidopsis, tobacco and rice are able to promote transcription in diverse systems (Rochon *et al.*, 2006; Maier *et al.*, 2011; Chern *et al.*, 2012), and it was suggested that NPR1 functions as transcription co-activator (Rochon *et al.*, 2006). NPR1 also interacts with the group of small NIM1 INTERACTING (NIMIN) proteins (Weigel *et al.*, 2001). Like *NPR1*, *NIMIN* genes are common in the whole higher plant kingdom. Arabidopsis contains four *NIMIN* genes. Of these, *NIMIN1* and *NIMIN2* are strongly up-regulated by SA (Weigel *et al.*, 2001, 2005; Glocova *et al.*, 2005; Hermann *et al.*, 2013). Similarly, tobacco *NIMIN2*-type mRNAs accumulate in response to the SA signal (Horvath *et al.*, 1998; Zwicker *et al.*, 2007). NIMIN1 (N1) and NIMIN2 (N2) possess similar motifs by which they bind to a domain identified in the NPR1 C-terminal third, termed N1/N2 binding domain (BD; Maier *et al.*, 2011). The functional significance of NIMIN proteins for NPR1 activity has been addressed in overexpression experiments. Both Arabidopsis NIMIN1 and NEGATIVE RGULATOR OF RESISTANCE (NRR), a NIMIN homolog from rice, are able to inhibit induction of *PR* genes and cause enhanced susceptibility to bacterial pathogens in plants (Weigel *et al.*, 2005; Chern *et al.*, 2005, 2008). Thus, NIMIN proteins modulate NPR1 activity.

The Arabidopsis NPR family comprises six members. NPR1 to NPR4 are closely related to each other sharing the same domain structure. All NPR1, NPR3 and NPR4 were reported to bind the SA signal molecule. Yet, the functional roles NPR proteins play with regard to perception and transduction of the defense hormone SA are discussed controversially. In a model proposed by Fu *et al.* (2012), NPR3 and NPR4 function as adaptors in cullin3-based E3 ligases. NPR3 and NPR4 interact with NPR1 in an SA-dependent manner thus controlling NPR1 levels during SAR through proteasomal degradation. While NPR3 and NPR4 were found to bind SA, NPR1 did not. More recently, Ding *et al.* (2018) have suggested that NPR3 and NPR4 act redundantly as transcriptional co-repressors of SA-responsive genes independently from NPR1, which clearly is a positive regulator of defense. All NPR3, NPR4 and NPR1 exert their function through binding SA. In another model, Wu *et al.* (2012) likewise claimed NPR1 an SA receptor. SA is bound via a metal ion in complex with two cysteine residues in the NPR1 C-terminus. Binding of SA activates tranquilized NPR1 leading to intramolecular rearrangements and transcription activity mediated via the NPR1 C-terminus.

In tobacco, the NPR family comprises only three members. NtNPR1 and NtNIM1-like1, which is most closely related to AtNPR3/NPR4, share the domain structure with AtNPR1. First biochemical evidence that some NPR proteins can indeed perceive SA directly came from analyses of NtNPR1, NtNIM1-like1 and AtNPR1 in heterologous yeast systems in absence of other plant proteins (Maier *et al.*, 2011). NtNPR1 gains additional transcription activity, when SA is added to yeast growth medium. Furthermore, binding of NIMIN1 and NIMIN2-type proteins to NtNPR1, NtNIM1-like1 and AtNPR1 is inhibited in presence of SA. Of note, the C-terminal third of NtNPR1 is sufficient to respond to the SA signal. The data indicate that NPR family members are able to sense SA via their C-termini, and that they undergo alterations upon perception of SA. SA sensitivity in the heterologous yeast system was mapped to the arginine residue embedded in the conserved pentaamino acid motif LENRV in tobacco and Arabidopsis NPR1 (R431 in NtNPR1, R432 in AtNPR1; Maier *et al.*, 2011). A mutant in the LENRV motif, R432K (nim1-4), abolishes chemically induced *PR-1* gene expression and resistance to fungi in Arabidopsis plants (Ryals *et al.*, 1997). Thus, NPR1 proteins contain two distinct regions in their C-terminal thirds which are both involved in processing the SA signal, the signature LENRV, or a related motif, including the essential arginine residue, and the binding domain for SA-induced NIMIN1 and NIMIN2 proteins. Importantly, conclusions deduced from biochemical analyses of NPR proteins in yeast are corroborated by genetic evidence provided through an en masse in planta screen for Arabidopsis insensitive to the functional SA analog BTH (Canet *et al.*, 2010). In this screen, 24 mutants were identified harboring a single amino acid exchange in the *NPR1* gene. Sixteen mutants were found to be clustered in two distinct regions in the C-terminus, thus providing structural insights into AtNPR1. One cluster corresponds to the LENRV signature and adjacent amino acids. The nim1-4 mutant was found three times in the screen. The second cluster is nearly identical to the N1/N2BD identified through biochemical dissection of tobacco and Arabidopsis NPR1 proteins in yeast. Overall, the findings strongly suggest that both the LENRV harboring region with the critical arginine residue and the N1/N2BD are intimately associated with the SA response.

To understand SA signal perception and transduction through NPR family proteins, we extended biochemical analysis of tobacco NPR1. NtNPR1 was chosen because it supports two distinct SA-dependent reactions in yeast. Here, we confirm that the plant immune signal SA acts directly on tobacco NPR1. We demonstrate physical interaction in yeast, in vitro, in plant and in animal cells between the two C-terminal regions LENRV and N1/N2 BD. In yeast, association of the two domains occurs spontaneously, but is enhanced considerably by SA depending on R431 and on intact LENRV and N1/N2BD interfaces. Importantly, physical association of LENRV and N1/N2BD produces transcription activity.

## Materials and Methods

### Yeast hybrid analyses

Yeast hybrid protein assays (Y1H, Y2H, Y3H) were conducted as reported earlier (Weigel *et al.*, 2001; Maier *et al.*, 2011). Yeast cells were grown in absence or presence of different chemicals as indicated. Compounds insoluble in water were solved in dimethyl sulfoxide (DMSO). The final DMSO concentration in yeast growth medium was 0.5%. *LacZ* reporter gene activities were tested in duplicate with at least three independent colonies. Experiments were repeated at least once with new yeast transformations. Representative results collected in one experimental set are shown. Results are depicted as mean enzyme activities in Miller units, plus and minus standard deviation (SD).

### In vitro pull-down assays

*NtNPR1* parts comprising LENRV (aa 386-465) and LENRV R431K were overexpressed as maltose binding protein (MBP) fusions in *E. coli*, and *NtNPR1(466-588)* encoding N1/N2BD was overexpressed in fusion with glutathione S-transferase (GST). Roughly equal molar amounts of fusion proteins (4μg MBP–NtNPR1(386-465) and 2μg GST–NtNPR1(466-588)) were incubated for 30 min at room temperature in PBS supplemented with 10% dry milk and 1% TritonX-100 in a total volume of 100μl. SA or non-functional analog 4-OH BA were added to binding reactions at a final concentration of 1mM, and CuSO_4_ was added at a final concentration of 50μM. After incubation, 100μl of glutathione agarose resin, equilibrated with PBS/dry milk/Triton, were added, and reactions were rotated for 1hr at room temperature. Bound proteins were collected by centrifugation, and pellets were washed twice with cold PBS. Pulled-down proteins were eluted from the resin by boiling in SDS sample buffer. Equal volumes of eluted proteins and of supernatants from binding reactions were subjected to SDS gel electrophoresis. GST–NtNPR1(466-588) proved to be unstable under binding conditions. To demonstrate equal distribution of the GST fusion protein in binding reactions, aliquots were immediately taken from assays prior to incubation and denatured in SDS sample buffer. Immunodetection of proteins was with antibodies against MBP and GST.

### Transient expression of *NtNPR1–Venus* half genes in *N. benthamiana* and in HEK293 cells

Transient gene expression by agroinfiltration in *Nicotiana benthamiana* was performed as described previously (Hermann *et al.*, 2013). Briefly, *NtNPR1–Venus* half gene constructs in pSPYNE2 or pSPYCE2 vectors (Nagai *et al.*, 2002; Walter *et al.*, 2004) were transferred to *Agrobacterium tumefaciens* strain GV3101 by triparental mating. Each bacterial strain was adjusted to an optical density (OD600) of 0.8 in 10mM MgCl2 supplemented with 150μM acetosyringone. For co-infiltration experiments, equal volumes of two bacterial suspensions were mixed with 1 volume of a strain carrying the p19 suppressor from *Tomato bushy stunt virus* (1+1+1). Mixtures of Agrobacterium suspensions were infiltrated in leaves of three different plants. One day before microscopic observation, plants were sprayed with water or 5mM SA. Epidermal peels were viewed under epifluorescence and bright field conditions 4 to 7 days post infiltration (dpi) as reported previously (Maier *et al.*, 2011). Proteins were extracted from leaf tissue exhibiting fluorescence with GUS lysis buffer, and equal volumes of crude extracts were loaded on SDS polyacrylamide gels. Immunodetection was performed on separate blots with an antiserum directed against GFP, with α-GST–NtNPR1(386-588) and with an antiserum raised against purified tobacco PR-1a protein (Stos-Zweifel *et al.*, 2018). Unspecific bands reacting with the antisera are marked to demonstrate equal gel loading.

Human embryonic kidney (HEK293) cells were maintained in DMEM/Ham’s medium supplemented with fetal bovine serum (10% FBS) and 5% penicillin/streptomycin at 37°C in a CO_2_ (5%) incubator. 5×10^5^ cells were transfected by the calcium phosphate method at a confluence of 70-80%. Two hours before transfection, growth medium was removed and replaced with fresh medium. For each transfection, 4μg DNA/construct were diluted in 100μl 0.3M CaCl_2_. The DNA solution was added dropwise to 100μl 2x HBS (Hepes buffered saline) while mixing gently. The mixture was incubated for 30 min at room temperature and then distributed evenly to one well of a six-well plate. SA was added to medium at a final concentration of 1mM after 18 hours of incubation. Cells were viewed under epifluorescence and bright field conditions 6.5hr after application of SA.

## Results

### Mutations in both the LENRV and N1/N2BD regions affect salicylic acid responsiveness of tobacco NPR1

The en masse in planta screen for Arabidopsis insensitive to BTH together with biochemical dissection of NPR proteins in yeast suggested that SA responsiveness of the SAR regulator NPR1 may rely not only on the arginine residue in the LENRV motif, but possibly also on the extended LENRV region and the binding domain for SA-induced NIMIN1 and NIMIN2 proteins (Fig. S1a,b). We exploited knowledge gained from the screen by introducing missense mutations identified in Arabidopsis *npr1* alleles into tobacco *NPR1* for biochemical analysis of mutant proteins in yeast one-hybrid (Y1H) and yeast two hybrid (Y2H) assays. Altogether, we generated four point mutations of *NtNPR1* at amino acid positions identical between Arabidopsis and tobacco NPR1 (Fig. S1c). Mutations were tested in the context of the NtNPR1 C-terminal third from amino acids 386 to 588. Two mutants, E442K and A450V, are associated with the LENRV region. The other two mutants, R491K and L495F, lie at the N-terminal boundary of the N1/N2BD (Figs 1a, S1c). As controls, we used mutants R431K (corresponding to nim1-4) and F505/506S, which lack SA sensitivity and N1/N2 binding potential, respectively (Maier *et al.*, 2011).

**Fig. 1.**
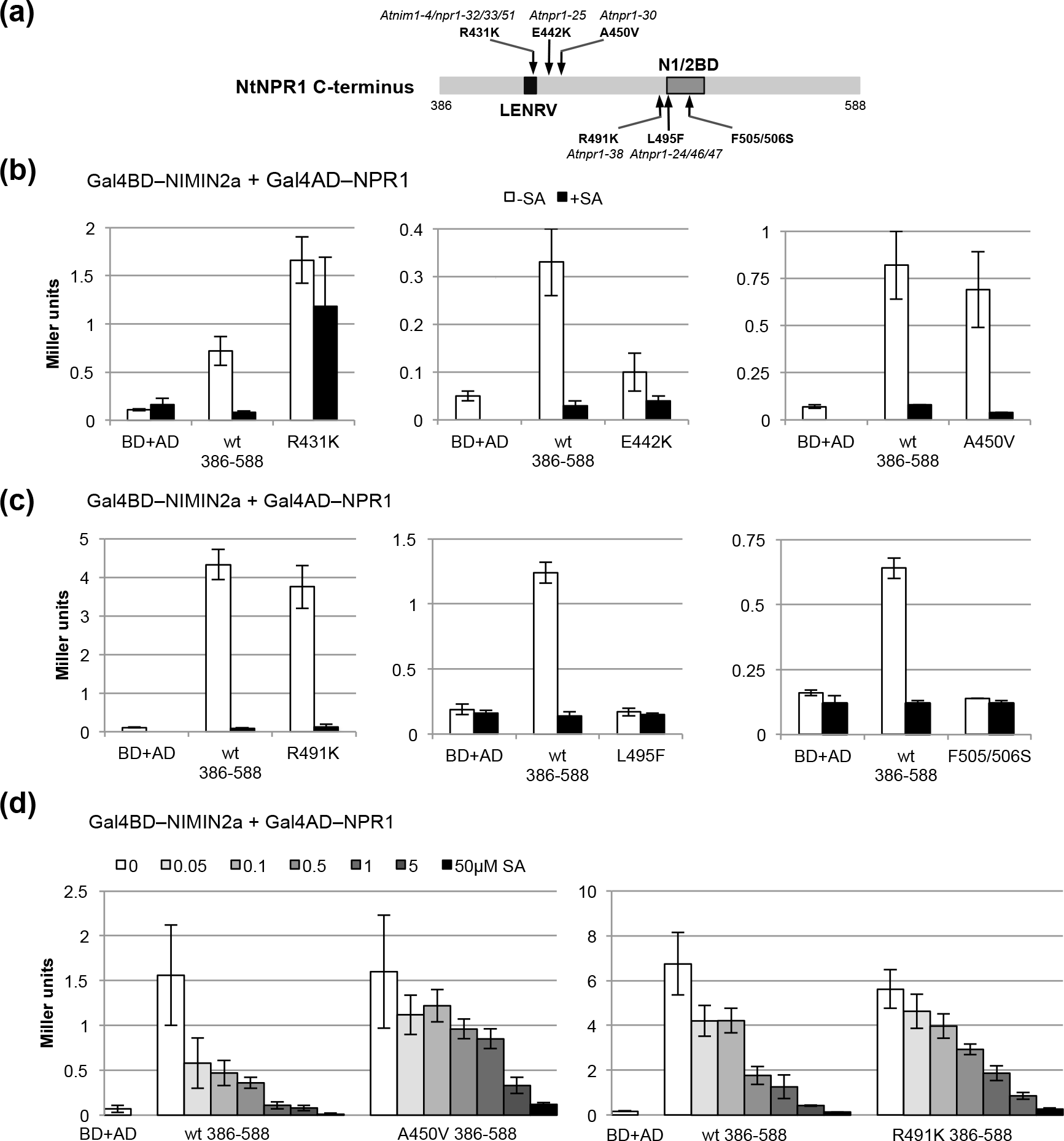
Interaction of tobacco NIMIN2a with the tobacco NPR1 C-terminal third is compromised by mutations in LENRV and N1/N2BD regions. (a) Domain structure of the tobacco NPR1 C-terminus and positions of mutations tested. (b,c) Interaction of NIMIN2a with tobacco NPR1 C-terminal proteins mutated in LENRV (b) or N1/N2BD (c) regions. Protein-protein interaction of mutant proteins was compared directly to interaction of wild-type NtNPR1(386-588) in quantitative Y2H assays. Interaction studies were performed in absence or presence of 100μM (b) or 300μM SA (c). (d) Salicylic acid dose response curves of interaction of NIMIN2a with tobacco NPR1 C-terminal proteins carrying mutations A450V or R491K. Protein-protein interaction of mutant proteins was compared directly to interaction of wild-type NtNPR1(386-588) in quantitative Y2H assays.

Initially, we tested interaction of mutant proteins with tobacco NIMIN2a (Fig. 1b,c). In accord with previous findings, mutant NPR1(386-588)F505/506S does not support NIMIN2a binding. Furthermore, mutants E442K and L495F are clearly compromised in NIMIN2a binding, while mutants R431K, A450V and R491K exhibit interaction similar to the wild-type. In presence of high SA levels, interaction of mutant proteins with NIMIN2a is inhibited with exception of the SA insensitive mutant R431K. Significantly, dose-response studies revealed that, although displaying normal NIMIN2a binding in absence of SA and full relief in presence of high SA, mutants A450V and R491K are nevertheless slightly affected in their overall SA response (Fig. 1d).

We also tested mutants with respect to their ability to support transcription of the *lacZ* reporter in Y1H assays (Fig. 2a,b). Mutant R431K is inactive irrespective whether tested with or without SA. Yet, the other mutants promote SA-induced l*acZ* reporter activity albeit to different extents. Dose-response relationships show that the transcription activation profile is shifted to higher SA concentrations with mutant proteins. The only exemption seems mutant R491K which displays a wild-type-like dose-response curve. Mutant NPR1(386-588)F505/506S is of particular interest. The mutant protein is clearly less sensitve to SA. On the other hand, the transcription activation potential is considerably enhanced at high SA concentrations as compared to the wild-type protein. The same effect is observed with the mutant full-length protein (Fig. 2b,c). In conclusion, SA sensitivity of NtNPR1 strictly depends on R431. Moreover, SA sensitivity of the NtNPR1 C-terminal third is affected, but not abolished, by different amino acid substitutions in both the extended LENRV and N1/N2BD regions.

**Fig. 2.**
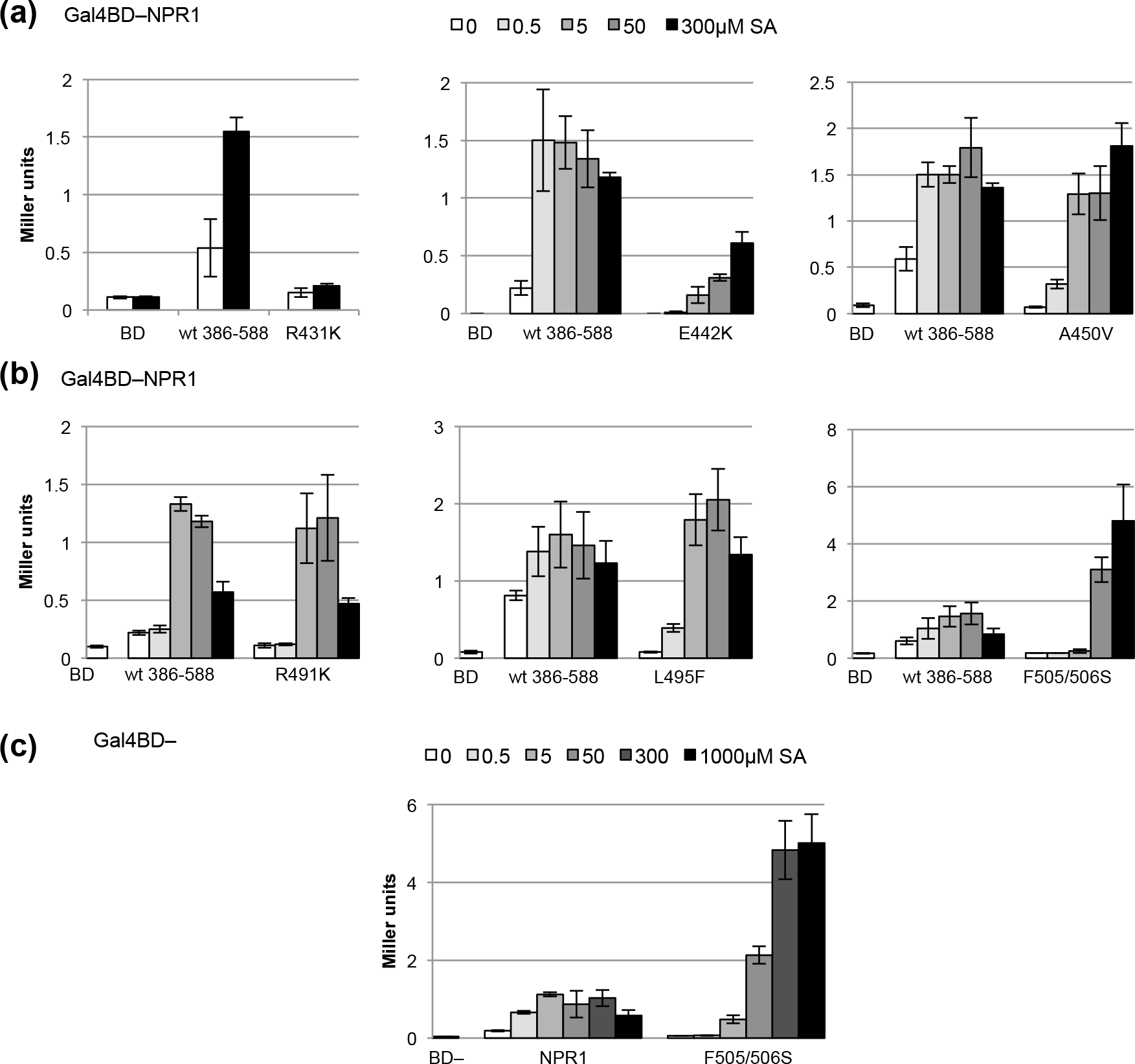
Transcriptional activity of the C-terminal third of tobacco NPR1 is compromised by mutations in LENRV and N1/N2BD regions. (a,b) Transcriptional activity of tobacco NPR1 C-terminal proteins mutated in LENRV (a) or N1/N2BD (b) regions. Transactivator activity of mutant proteins was compared directly to wild-type Gal4BD–NtNPR1(386-588) in quantitative Y1H assays. Assays were conducted in absence or presence of salicylic acid (300μM) or as dose response relationships. (c) Transcriptional activity of full-length tobacco NPR1 carrying mutations F505/506S. Transactivator activity of the mutant protein was compared directly to wild-type Gal4BD–NtNPR1 in quantitative Y1H assays. Salicylic acid dose response relationships are shown.

### Salicylic acid promotes physical association between LENRV and N1/N2BD parts of tobacco NPR1

Our findings suggested that the two C-terminal domains LENRV and N1/N2BD of tobacco NPR1 may co-operate in perception and/or transduction of the SA signal acting in concert with R431. To test whether the two domains may engage in physical interaction with each other, we separated the LENRV region from the N1/N2BD in the middle of the intervening sequence (Fig. 3a). The resulting parts, NtNPR1(386-465) and NtNPR1(466-588), were analyzed for protein-protein interaction potential in absence and presence of SA.

**Fig. 3.**
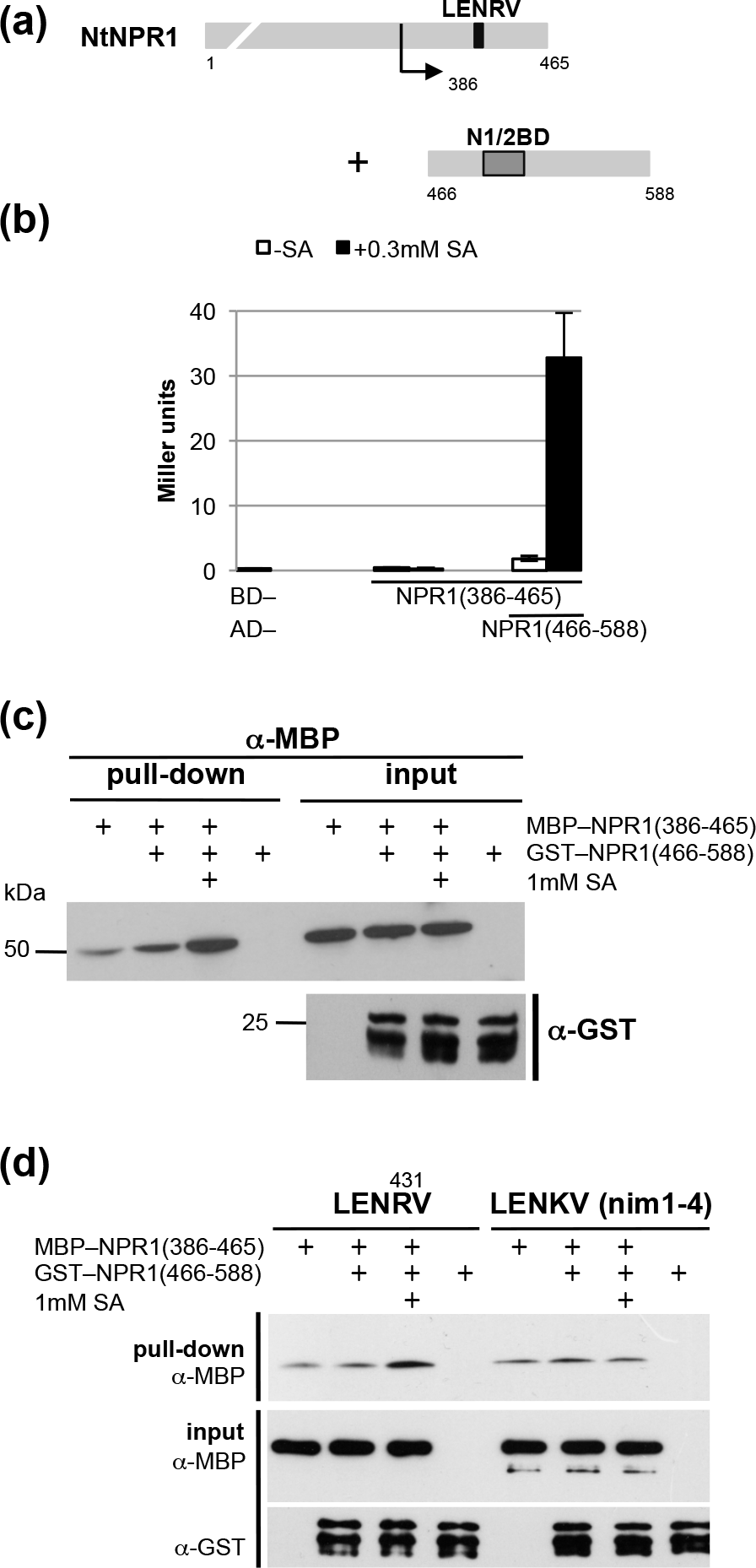
LENRV and N1/N2BD regions of tobacco NPR1 interact physically. (a) Separation of the NtNPR1 C-terminal region in the middle of the intervening sequence between the LENRV motif and the N1/N2BD. (b) Interaction of LENRV and N1/N2BD parts of NtNPR1 in yeast. Protein-protein interaction in absence or presence of SA was determined in quantitative Y2H assays. (c) In vitro pull-down assay of MBP–NtNPR1(386-465) with GST–NtNPR1(466-588). Binding reactions were performed in absence and presence of SA. Proteins were pulled down with glutathione sepharose and detected with an antiserum directed against maltose binding protein (MBP). Aliquots of binding reactions were detected with an antiserum directed against MBP after incubation and with GST antiserum before incubation. (d) In vitro pull-down assay of mutant protein MBP–NtNPR1(386-465) R431K with GST–NtNPR1(466-588).

Although NPR1(386-588) harbors transcription activity and binds NIMIN proteins in yeast, we were not able to determine such activities for the newly generated NPR1 deletions in Y1H and Y2H tests (Figs 3b, S2). Quite surprisingly, however, co-expression of fusion genes for Gal4 DNA-binding domain (GBD)–NPR1(386-465) and Gal4 transcription activating domain (GAD)–NPR1(466-588) yielded low level *lacZ* activity above the background in Y2H assays, indeed revealing loose spontaneous contact between the extended LENRV region and the N1/N2BD containing deletion (Fig. 3b). Interaction is enhanced largely by addition of SA to yeast culture medium. Furthermore, SA-dependent interaction between LENRV and N1/N2BD parts occurs also with the reciprocal Gal4 fusion proteins, with a N1/N2BD protein truncated at the C-terminus, GAD–NPR1(466-525), and in the full-length context when GBD–NPR1(1-465) is provided as partner for GAD–NPR1(466-588) (Fig. S3).

To verify direct contact between LENRV and N1/N2BD regions of tobacco NPR1 by an independent approach, we performed in vitro pull-down assays. *NPR1(386-465)* was expressed in *E. coli* fused to the sequence for maltose binding protein (MBP), and *NPR1(466-588)* was expressed in fusion with the sequence for glutathione S-transferase (GST). Binding reactions were incubated with glutathione resin, and pulled-down MBP–NPR1(386-465) was detected with an antiserum directed against MBP. We also tested aliquots of the binding reactions for amounts of input proteins. While MBP–NPR1(386-465) was readily detected as intact band in the supernatants of binding reactions, GST–NPR1(466-588) proved unstable (Fig. 3c,d and data not shown). Consequently, pull-down assays produced rather faint bands only. We found weak unspecific binding of MBP–NPR1(386-465) to the glutathione matrix (Figs 3c,d, S4). Typically, the background signal was slightly intensified in presence of GST–NPR1(466-588) possibly indicating spontaneous association between the two NPR1 parts. Clearly, addition of SA to binding assays reinforced interaction of MBP–NPR1(386-465) with GST–NPR1(466-588), while addition of Cu2+ ions had no effect (Figs 3c,d, S4). Furthermore, mutant protein MBP– NPR1(386–465)R431K, carrying the nim1-4 mutation, does not support SA-dependent interaction with GST–NPR1(466-588) (Fig. 3d).

We also tested whether the split C-terminal tobacco NPR1 regions could associate in plant cells using bimolecular fluorescence complementation (BiFC). The sequence harboring the LENRV motif was fused to the N-terminal part of *Venus (NPR1(386-465)–nVenus)*, and the N1/N2BD containing sequence was fused to the C-terminal part of *Venus (NPR1(466-588)– cVenus)*. Both fusion genes were expressed from the *CaMV35S* promoter. The constructs were transferred to *Agrobacterium tumefaciens*, and Venus complementation was studied after agroinfiltration in *N. benthamiana* leaves by epifluorescence microscopy. Co-expression of *Venus* half genes did not yield significant fluorescence, while co-expression of *NPR1(386-465)–nVenus* and *NPR1(466-588)–cVenus* produced strong fluorescence in the nucleus and in the cytosol of epidermal cells (Fig. 4a). Similar results were obtained with intact Venus protein. Accumulation of NPR1(386-465)–nVenus was demonstrated by immunodetection using antisera raised against GFP and GST–NtNPR1(386–588), respectively (Fig. S5a). However, accumulation of NPR1(466-588)–cVenus could not be shown explicitly with these antisera. In some cases, leaf tissue exposed to SA displayed enhanced fluorescence signals (Fig. 4a), but in most cases, we were not able to detect clear differences between water and SA-treated tissue (Fig. S5c,d). This may be due to uneven expression of fusion genes in different plants, to the more qualitative nature of fluorescence images or to the fact that the endogenous SA response is induced to some extent by the agroinfiltration procedure. Indeed, we found accumulation of PR-1 proteins to varying levels in agroinfiltrated *N. benthamiana* leaves expressing *Venus* half genes (Fig. S5b and data not shown). Fluorescence complementation was also observed with the reciprocal *Venus* half gene fusions (Fig. S5d).

**Fig. 4.**
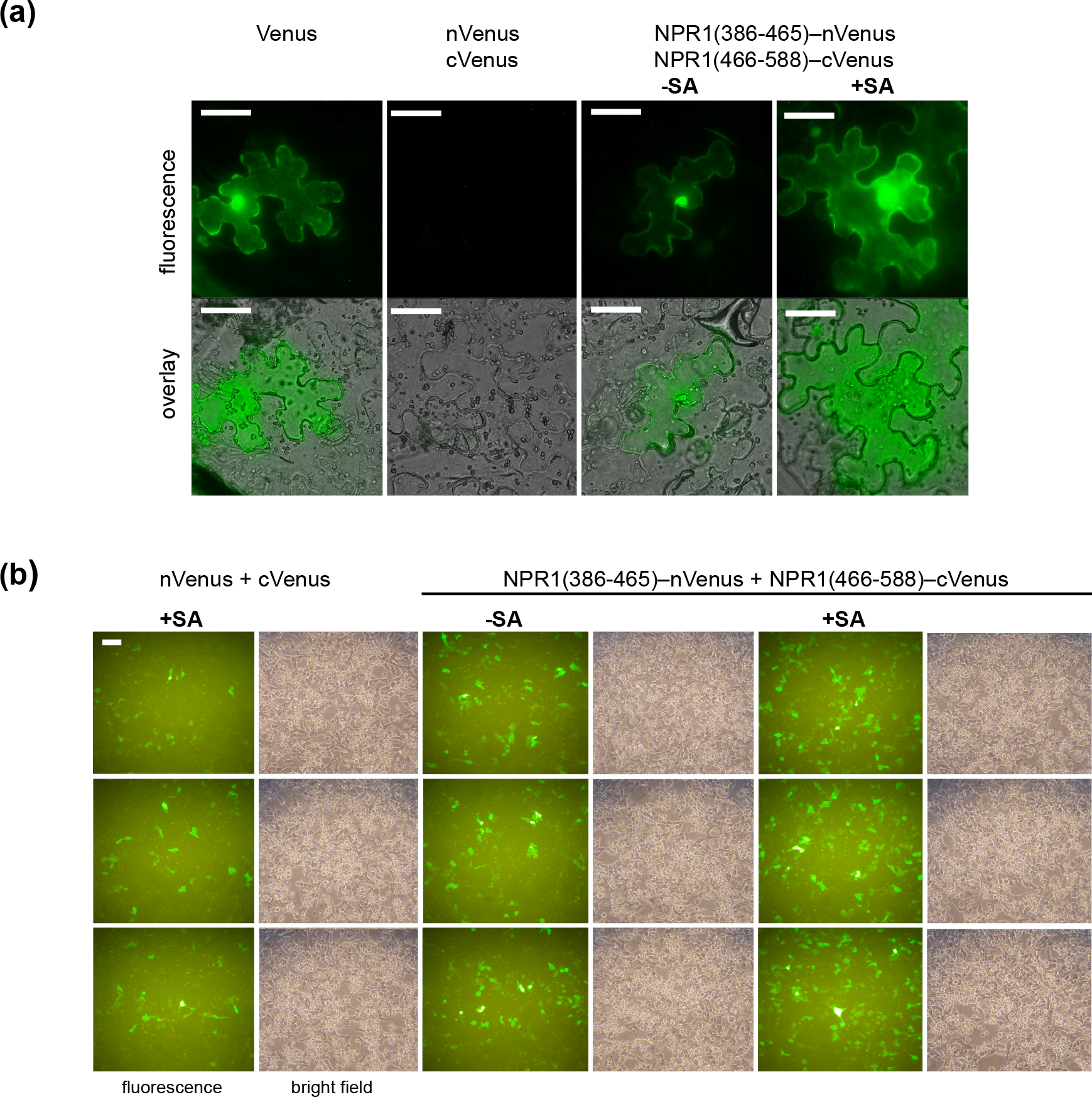
Visualization of LENRV and N1/N2BD interaction of tobacco NPR1 in plant and animal cells by bimolecular fluorescence complementation (BiFC). Cells were viewed under epifluorescence and bright field conditions. Representative results are shown. (a) LENRV–N1/N2BD interaction in plant cells. Mixtures of Agrobacterium suspensions harboring gene constructs as indicated were infiltrated in *N. benthamiana* leaf tissue. Epidermal peels were viewed 5 days post infiltration. One day prior to microscopy, plants were sprayed with water or 5mM SA. Scale bars, 50μm. (b) LENRV–N1/N2BD interaction in animal cells. HEK293 cells were transfected with mixtures of plasmid DNAs as indicated. Cells were viewed 1 day post transfection and 6.5hr after addition of SA to medium (final concentration 1mM). Results from three independent transfection experiments are shown. Scale bar, 200μm.

To further test the potential of tobacco NPR1 LENRV and N1/N2BD parts to serve as SA sensor, we expressed *NtNPR1–Venus* half gene fusions under control of the immediate early CMV promoter in HEK293 cells. We observed some background fluorescence in cells expressing *nVenus* and *cVenus* only. However, co-expression of *NPR1(386-465)–nVenus* and *NPR1(466-588)–cVenus* clearly enhanced fluorescence signals, i.e., by far more cells display fluorescence, and fluorescence in individual cells is generally stronger (Fig. 4b). The signals were even more intensified when the medium was supplemented with SA. Together, the data substantiate our hypothesis of SA-mediated intramolecular binding occurring between LENRV and N1/N2BD parts of tobacco NPR1.

### Functional salicylic acid analogs promote physical association between LENRV and N1/N2BD parts of tobacco NPR1

To underpin the relevance of contact between the LENRV region and the N1/N2BD for induction of SAR genes in planta, we used various SA analogs in the split NtNPR1 C-terminus Y2H assay. 2,6-Dichloroisonicotinic acid (INA) and benzothiadiazole (BTH) are chemical inducers of SAR and *PR-1* gene expression (Métraux *et al.*, 1991; Friedrich *et al.*, 1996; Görlach *et al.*, 1996). The compounds are structurally related to SA (Fig. S6a). In accord with their roles as functional SA analogs, all are able to promote association between GBD–NPR1(386-465) and GAD–NPR1(466-588) in a dose-dependent manner in yeast with BTH free acid being the most potent chemical (Fig. S6b). While INA and BTH can replace SA as signaling molecules in SAR, compounds like azelaic acid, β-aminobutyric acid (BABA) and pipecolic acid are implicated in spreading and amplifying the infection signal via unknown mechanisms acting on SA signaling (Jung *et al.*, 2009; Zimmerli *et al.*, 2000; Navarova *et al.*, 2012). Consistently, these compounds, which are structurally dissimilar to SA, remained inactive in the Y2H association assay (Fig. S7).

Previously, we have shown that SA sensitivity of tobacco NPR1 in yeast is imitated by benzoic acid (BA), an analog functional in resistance and *PR-1* gene induction in planta, but not by the non-functional analog 4-hydroxy benzoic acid (4-OH BA; White, 1979; Maier *et al.*, 2011). BA, similar to SA, is able to promote association, while 4-OH BA could not enhance weak spontaneous binding between LENRV and N1/N2BD parts in yeast (Fig. S8b). Likewise, 4-OH BA does not support interaction between MBP–NPR1(386-465) and GST–NPR1(466-588) in vitro (Fig. S4). In addition, we tested other BA derivatives known to function as inducers of defense in Arabidopsis (Fig. S8a; Knoth *et al.*, 2009). Chlorinated BA derivatives generally inhibited yeast cell growth when added to culture medium at higher doses. Therefore, we used the compunds only up to concentrations of 30μM that do not show effects. Anthranilic acid (AA) is clearly less active than BA, while 3,5-dichloroanthranilic acid (3,5-DCA) turned out to be the most active compound tested, even more potent than SA (Fig. S8c). We used SA and 3,5-DCA to compare the different SA-dependent reactions of tobacco NPR1 in yeast, i.e., association of C-terminal LENRV and N1/N2BD parts as well as gain of transcription activity and inhibition of NIMIN2 binding in full-length NtNPR1 context. Half maximal effective concentrations are similar for both chemicals and for all activities tested ranging from 3.2μM to 0.3μM (Fig. 5). The data indicate that 3,5-DCA is likely to act on NPR1 in a similar manner as SA.

**Fig. 5.**
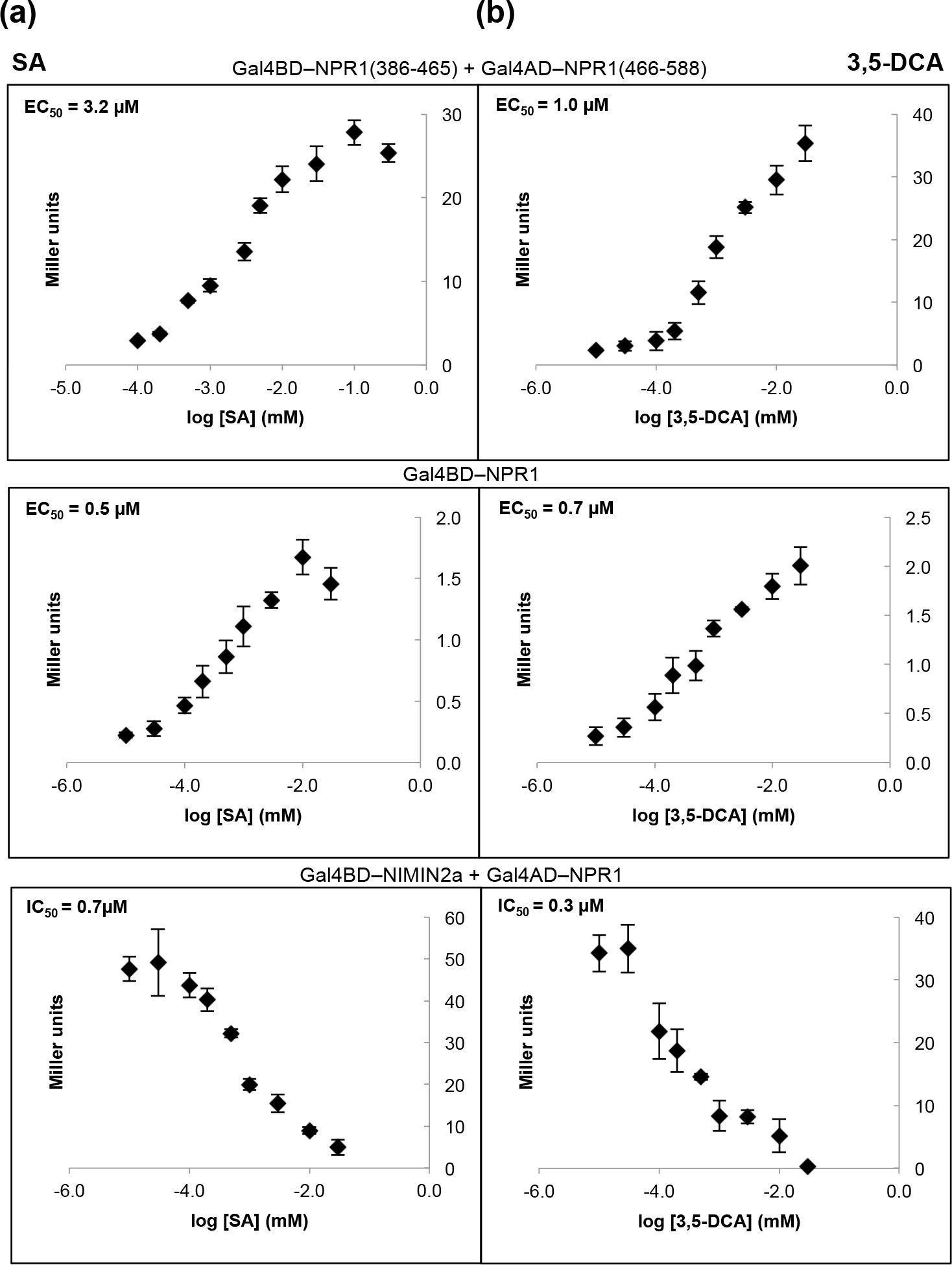
Salicylic acid-induced activities are coordinated in tobacco NPR1. (a) Estimation of half maximal effective concentration values of salicylic acid for association of LENRV and N1/N2BD parts, for transcription activation of NtNPR1 and for inhibition of NIMIN2a binding. Effects of different concentrations of SA (0.01 to 300μM) on NtNPR1 activities were determined in Y2H and Y1H assays as indicated. (b) Estimation of half maximal effective concentration values of 3,5-dichloroanthranilic acid (3,5-DCA) for association of LENRV and N1/N2BD parts, for transcription activation of NtNPR1 and for inhibition of NIMIN2a binding. Effects of different concentrations of 3,5-DCA (0.01 to 30μM) on NtNPR1 activities were determined in Y2H and Y1H assays as indicated. 3,5-DCA was solved in DMSO. The final DMSO concentration in yeast growth medium was 0.5%.

### Mutations in both the LENRV and N1/N2BD regions compromise their salicylic acid-dependent association

We showed that missense mutations, mapped in Arabidopsis *npr1* alleles in planta and introduced in LENRV and N1/N2BD regions of tobacco *NPR1*, impair SA sensitivity of the NtNPR1 C-terminal third. Furthermore, we demonstrated that SA-dependent association of LENRV and N1/N2BD parts occurs coordinately with both SA-dependent transcription activation of NtNPR1 and relief from NtNIMIN2a. Consequently, we would expect that mutations in the C-terminus likewise affect physical association of GBD–NPR1(386-465) and GAD–NPR1(466-588).

With mutant R431K, we observed neither spontaneous nor SA-induced association of C-terminal domains in Y2H, and with mutant F505/506S SA-dependent interaction was drastically reduced, although it did not seem completely abolished (Fig. 6a,b). Both mutant fusion proteins accumulate in yeast (Fig. 6c,d). The other mutants were compromised to different extents in the split NPR1 C-terminus Y2H assay (Fig. 6a,b). Mutant R491K was only slightly affected, while mutants E442K and L495F, located more closely to R431 and the N1/N2BD core, respectively, displayed the most severe effects. We also analyzed another protein mutated in the critical arginine residue, R431F (Fig. S9a). Mutant NPR1(386-588)R431F binds NIMIN2 proteins, but binding is insensitive to SA (Fig. S9b). Moreover, like mutant R431K, mutant R431F is neither able to induce transcription activity in the C-terminal region nor to associate LENRV R431F and N1/N2BD parts in an SA-dependent fashion (Fig. S9c,d). Together, the data suggest that the LENRV and N1/N2BD regions of tobacco NPR1 contact each other. Contact is driven by SA and strictly relies on R431. Furthermore, effective association requires intact LENRV and N1/N2BD interfaces.

**Fig. 6.**
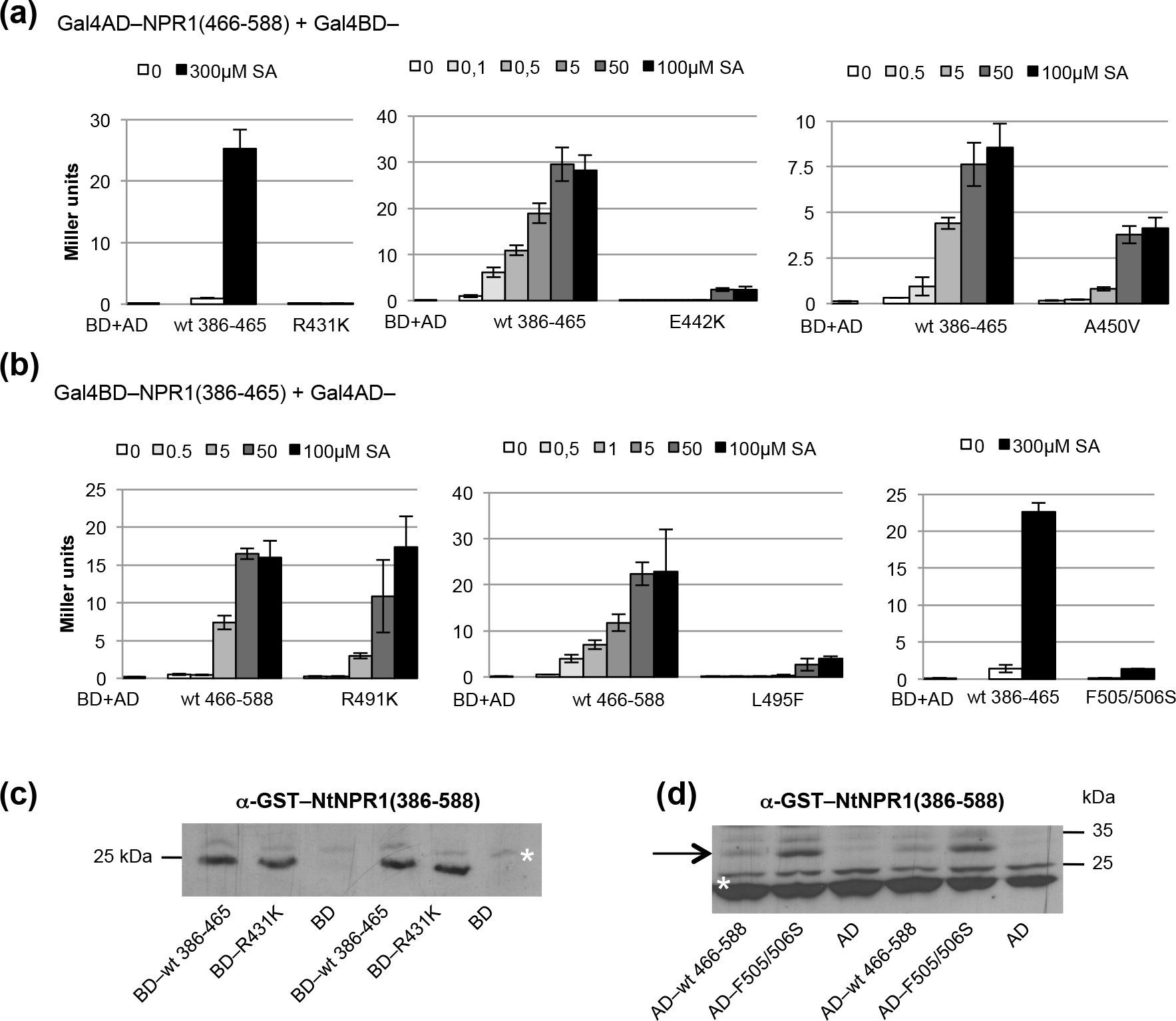
Mutations in LENRV and N1/N2BD regions of tobacco NPR1 impair their physical association in yeast. (a,b) Interaction of tobacco NPR1 C-terminal proteins mutated in LENRV (a) or N1/N2BD (b) regions. Interaction of mutant proteins was compared directly to interaction of wild-type LENRV and N1/N2BD parts in quantitative Y2H assays. Assays were conducted in absence or presence of salicylic acid (300μM) or as dose response relationships. (c,d) Accumulation of mutant proteins GBD–NtNPR1(386-465)R431K (c) and GAD–NtNPR1(466-588)F505/506S (d) in yeast. Protein extracts from two independent colonies each transformed with empty vector or with vectors encoding the respective wild-type or mutant proteins were analyzed by immunodetection with an antiserum directed against GST–NtNPR1(386-588). The position of GAD–NtNPR1(466-588) in the gel is marked by an arrow, and unspecific bands indicating equal loading of the gels are marked by asterisks.

### Salicylic acid-mediated interaction between LENRV and N1/N2BD regions of tobacco NPR1 produces transcription activity

Next, we addressed whether LENRV and N1/N2BD interaction promoted by SA would produce activity in the tobacco NPR1 C-terminus. In particular, we asked whether association of LENRV and N1/N2BD would result in transcription activity. This speculation appeared plausible as the NtNPR1 C-terminus fused to GBD mediates low level transcription activity in yeast that is clearly enhanced in presence of SA.

The seqences encoding NtNPR1 C-terminal parts were cloned in a yeast tri-hybrid (Y3H) vector. The sequence for the LENRV region from amino acids 386 to 465 was expressed fused to the sequence for GBD, and the sequence for N1/N2BD (aa 466-588) was put under control of the *MET25* promoter which is repressed in presence and de-repressed in absence of methionine (Fig. 7a; Tirode *et al.*, 1997). Without addition of SA, we did not detect any reporter activity (Fig. 7b). However, when yeast growth medium was supplemented with SA, clear *lacZ* activity was observed. This activity is reduced when N1/N2BD accumulation is repressed by addition of methionine to yeast culture medium. With mutant proteins LENRV R431K and N1/N2BD F505/506S we did not detect transcription activity (Figs 7c, S10a). However, mutant N1/N2BD R491K, supporting SA-mediated association of C-terminal LENRV and N1/N2BD R491K parts, yields wild-type-like transcription activity (Fig. S10b). Together, SA induces interaction between LENRV and N1/N2BD parts of tobacco NPR1 which, in turn, produces transcription activity in yeast.

**Fig. 7.**
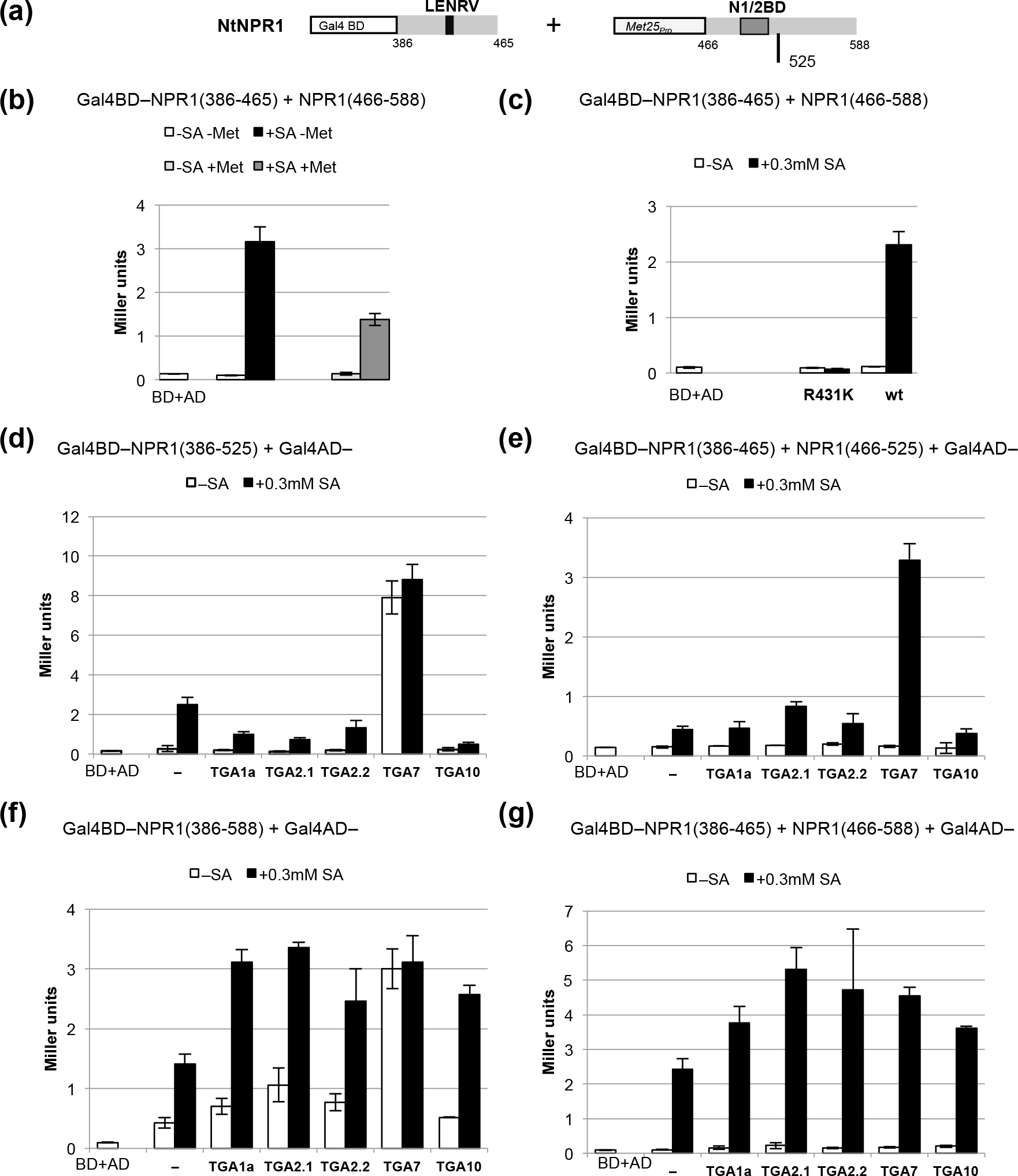
Association of LENRV and N1/N2BD parts of tobacco NPR1 produces transcription activity in yeast and enables TGA transcription factor interaction. (a) Design of yeast hybrid experiments shown in panels b and c. The sequence for the LENRV domain is fused to the sequence for Gal4-DNA binding domain (GBD), and the N1/N2BD encoding sequence is expressed from the *MET25* promoter. (b) Association of LENRV and N1/ N2BD parts by salicylic acid produces transcription activity. Quantitative assays were conducted with yeast cells grown without or with methionine and in absence or presence of salicylic acid (300μM). (c) Mutant R431K cannot reconstitute a transcriptionally active C-terminus. Yeast cells were grown in medium lacking methionine. Quantitative hybrid assays were performed with mutant GBD–NtNPR1(386-588)R431K and wild-type protein in absence or presence of salicylic acid. (d,f) Interaction of the C-terminal region of tobacco NPR1 with tobacco TGA factors. Quantitative Y2H assays were performed with deletion GBD-NtNPR1(386-525) (d) and with GBD–NtNPR1(386-588) (f) in absence and presence of salicylic acid. (e,g) Interaction of TGA factors with the tobacco NPR1 C-terminus reconstituted from LENRV and N1/N2BD parts by SA. Reconstitution experiments were conducted with LENRV(386-465) part of NtNPR1 and with N1/ N2BD truncated at amino acid 525 (e) or with N1/N2BD(386-588) (g).

### Salicylic acid-induced transactivating domain resides at the tobacco NPR1 C-terminus

The tobacco NPR1 C-terminus fused to GBD and the Y3H vector construct we used for reconstitution of the NtNPR1 C-terminus do not harbor plasmid-encoded transcription activation domains. Therefore, the SA-activated NtNPR1 C-terminus, itself, must be able to contact the yeast transcriptional machinery eliciting reporter gene expression. To map the site of transcription activation, we used NtNPR1 truncated at amino acid 525. Recently, we have isolated TGA7, a novel member of the tobacco TGA family, that interacts preferentially and uniquely with the NtNPR1 C-terminus. The factor binds constitutively around the N1/N2BD, independently from SA (Fig. 7d,f; Stos-Zweifel *et al.*, 2018). In contrast, NtTGA2.1 interacts with the N-terminal part of NtNPR1. Both TGA7 and TGA2.1 display transcription actitvity in yeast.

Importantly, factors TGA7 and TGA2.1 cannot bind to split LENRV and N1/N2BD regions (Fig. S11). Yet, TGA7 interacts with the C-terminus reconstituted by SA from LENRV and N1/N2BD(466-525) parts (Fig. 7e). Of note, the C-terminus truncated at position 525 produces only rather weak transcription activity in presence of SA (Figs 7d,e, S12). It interacts primarily with transcription factor TGA7. By contrast, the full C-terminus from positions 386 to 588, as a whole or reconstituted from LENRV and N1/N2BD parts, yields moderate transcription activity with SA and furthermore supports interaction with any tobacco TGA factor we tested, even with TGA1a and TGA10 classified as NPR1 non-interactors (Fig. 7f,g; Niggeweg *et al.*, 2000; Schiermeyer *et al.*, 2003). Thus, TGA factors can interact with the NtNPR1 C-terminus at distinct sites in an SA-dependent and SA-independent manner. While TGA7 binds constitutively in vicinity to the N1/N2BD, the last 63 amino acids of tobacco NPR1 mediate SA-induced contacts to the yeast transcription machinery which may imply TGA factor interactions.

### Salicylic acid inducible NIMIN proteins can bridge LENRV and N1/N2BD parts of tobacco NPR1

Biochemical dissection of tobacco NPR1 suggests that the binding domain for SA-induced NIMIN proteins is critical for transduction of the SA signal. Therefore, we would infer that NIMIN binding to the NPR1 C-terminus could affect SA signaling.

In fact, expression of *NtNIMIN2a* or *NtNIMIN2c* from the *MET25* promoter represses constitutive transcription activity of GBD–NtNPR1(386-588) in yeast (Fig. 8a). Upon addition of SA to medium, repression is partially relieved suggesting that NIMIN proteins can occlude LENRV–N1/N2BD interaction. To test if NIMINs could contact the LENRV region directly, we co-expressed *GBD-NtNPR1(386-465)*, *NtNPR1(466-588)* under control of the conditional *MET25* promoter and *GAD–NtNIMINs* in yeast (Fig. 8b). We indeed observed low level transcription activity in absence of methionine, which is reduced in presence of methionine, indicating that NtNIMIN2a and a tobacco NIMIN1-like protein can form complexes together with separated LENRV and N1/N2BD parts of NtNPR1 (Fig. 8c). Complex formation could also be demonstrated with mutant protein N1/N2BD R491K, but not with mutation R431K introduced into the LENRV part (Fig. S13). However, even minor concentrations of SA added to yeast growth medium (0.5μM) already favored direct contact between LENRV and N1/N2BD regions thus overshadowing the bridging effect of NIMIN1 and NIMIN2 proteins (data not shown).

**Fig. 8.**
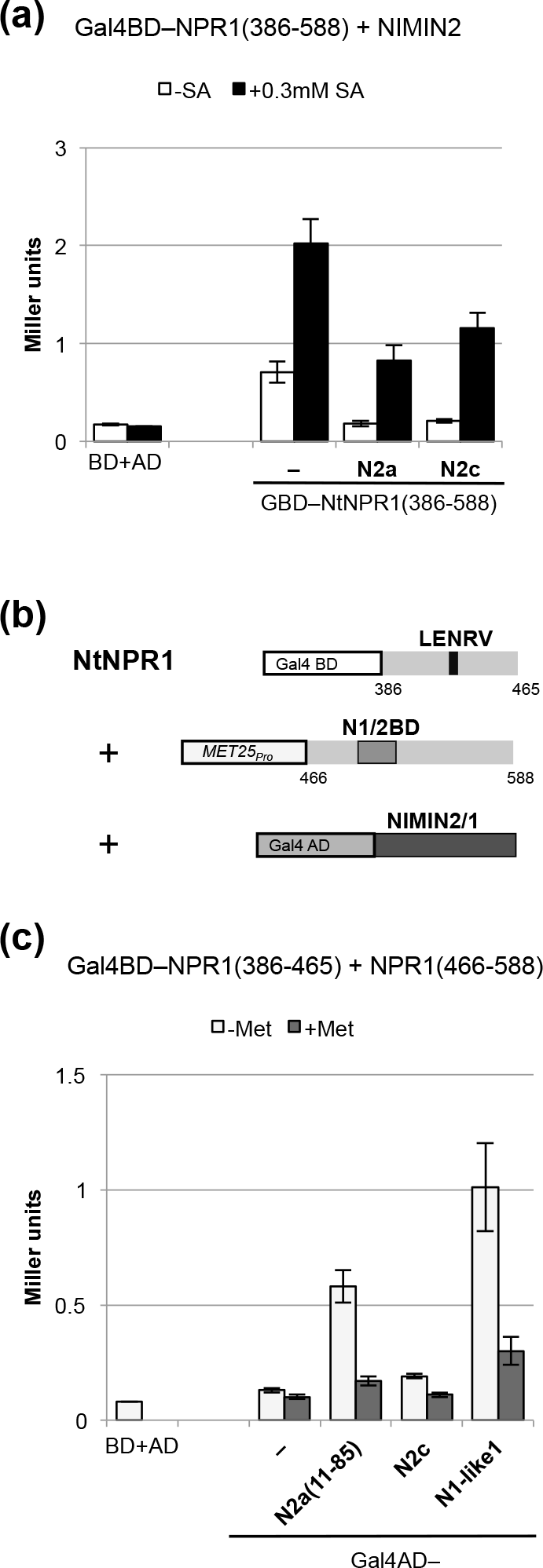
NIMIN proteins affect SA signaling through the tobacco NPR1 C-terminus. (a) Transcription activity of the tobacco NPR1 C-terminus is compromised in presence of NIMIN2 proteins. Transcription activity of GBD-NtNPR1(386-588) was determined in absence and presence of NtNIMIN2a or NtNIMIN2c in quantitative interaction assays in yeast. *NIMIN2* genes were expressed from the *MET25* promoter without addition of methionine to medium. (b) Scheme depicting the experiment shown in panel c. (c) NIMIN proteins can bridge LENRV and N1/N2BD parts of the tobacco NPR1 C-terminus. Quantitative Y3H interaction assays were performed with cells grown in absence or presence of methionine.

## Discussion

Resolving the mechanism of SA signaling through NPR1, the central regulator of SAR, has been a major focus during the past two decades. Different models have been launched which are, at least in parts, contradictory. While the NPR1–TGA factor connection, bringing NPR1 in proximity to promoter regions of SA-induced *PR-1* genes, is well established, the NPR1–NIMIN connection has been disregarded on the whole. NIMIN proteins bind strongly to NPR1, and interaction of SA-induced NIMIN1 and NIMIN2-type proteins occurs at a well-defined site in the C-terminal third of NPR1, referred to as N1/N2BD. Apart from binding NIMIN proteins, the C-terminus of NPR1 has also been recognized as direct target for the SA signal and as platform for interaction with transcription factors.

### Salicylic acid induces intramolecular interaction between LENRV and N1/N2BD regions that results in activity changes of tobacco NPR1

Stimulated by genetic and biochemical evidence, we found physical association between C-terminal LENRV and N1/N2BD parts of tobacco NPR1 in yeast, in vitro, in plant and in animal cells. In yeast, separated LENRV and N1/N2BD regions are able to associate spontaneously. This weak interaction is considerably enhanced by SA and functional analogs, but not by the non-functional analog 4-OH BA. Our data corroborate previous reports on in vitro binding of SA to Arabidopsis NPR1 (Wu *et al.*, 2012; Kuai *et al.*, 2015; Manohar *et al.*, 2015; Ding *et al.*, 2018). Most importantly, SA-mediated physical association of LENRV and N1/N2BD parts produces transcription activity in yeast. Altogether, the tobacco NPR1 C-terminal third supports three SA-dependent reactions, i.e., gain of transcription activity, relief from NIMIN1/NIMIN2 binding and intramolecular interaction of LENRV and N1/N2BD parts. These reactions occur coordinately. Presumably, SA urges remodeling in the NtNPR1 C-terminus involving LENRV and N1/N2BD regions, and remodeling, in turn, can initiate both expulsion of NIMIN proteins from N1/N2BD and activation of gene transcription.

### The arginine residue in the conserved LENRV motif of NPR1 is essential for perception of the SA signal

Although mutants in LENRV and N1/N2BD regions we tested were generally affected in their activities, the mutation mostly impacting tobacco NPR1 is R431K. The mutant protein is entirely insensitive to SA, but it is still able to bind NIMIN proteins. The corresponding mutant was isolated several times in genetic screens from Arabidopsis using two different screening regimes (Ryals *et al.*, 1997; Canet *et al.*, 2010). The arginine residue lies within the conserved pentaamino acid motif LENRV (Maier *et al.*, 2011). Notably, mutant protein AtNPR1 R432Q does not bind SA (Ding *et al.*, 2018). Collectively, the data suggest that the arginine residue in the conserved LENRV motif may be directly involved in perceiving the SA signal molecule. It seems conceivable that the positively charged guanidino group of arginine could complex the carboxyl of SA thus neutralizing a positive charge and jamming a hydrophobic benzene ring in the amino acid chain. As a consequence, local protein conformation could be disrupted allowing the benzene ring of SA to fish for contacts among side chains from nearby residues of the LENRV and/or N1/N2BD regions. Alternatively, disruption of local protein conformation by SA binding may be sufficient to build contacts among residues from LENRV and N1/N2BD regions. This, in turn, would shape a novel fold, molded by SA, that could enable, for example, exposure to the transcription machinery. Together, evidence provided by us and others strongly suggests that SA interacts directly with the arginine in the conserved LENRV motif to transduce the SA signal via rearrangement of the NPR1 C-terminus.

### Tobacco NPR1 harbors multiple docking sites for TGA transcription factors

By biochemical dissection in yeast, tobacco NPR1 reveals at least three distinct sites for interaction with members of the tobacco TGA transcription factor family. NtTGA2.1 and NtTGA2.2 contact NtNPR1 within the region from amino acids 1 to 465 (Stos-Zweifel *et al.*, 2018). Binding is constitutive, independent from SA. Similarly, the binding site for AtTGA2, AtTGA3 and AtTGA5 was mapped to the central ankyrin repeat domain of Arabidopsis NPR1 (Zhang *et al.*, 1999; Zhou *et al.*, 2000). NtTGA7, on the other hand, contacts tobacco NPR1 primarily near the N1/N2BD. Binding occurs constitutively, but is outcompeted by presence of NtNIMIN2 proteins (Stos-Zweifel *et al.*, 2018). In addition, tobacco NPR1 harbors an SA-inducible transcription activation domain at the C-terminus from positions 526 to 588. Transcription activity from this domain is enhanced by any tobacco TGA protein we tested implying that the domain can interact broadly with transcription factors. The SA-induced site of TGA factor interplay at the NtNPR1 C-terminus is distinct from the site of constitutive NtTGA7 interaction. Apparently, docking sites on NtNPR1 are used differentially by members of the TGA family and accessibility can change depending on physiological cues provoking rearrangements of the protein. In fact, NtNPR1 is the only NPR protein from tobacco and Arabidopsis, not including BOP proteins, that exhibits transcription activity in yeast. However, although AtNPR1 and the AtNPR1 C-terminus are inactive in yeast (Stos-Zweifel *et al.*, 2018), it has been shown by in vivo plant transcription assays that AtNPR1 harbors an autonomous transactivation domain in the last 80 residues (Rochon *et al.*, 2006; Wu *et al.*, 2012). Thus, functional similarities among NPR1 proteins from tobacco and Arabidopsis appear to pertain both perception of SA through the arginine in the LENRV motif and transduction of the SA signal.

### NIMIN1 and NIMIN2 proteins shape the NPR1 C-terminus

In this scenario, it is of interest that SA-induced NIMIN1 and NIMIN2 proteins bind to NPR1 close to the site of SA perception. In NtNPR1, the N1/N2BD itself, by physical interaction with the LENRV region, is involved in transducing the SA signal. Most importantly, NIMIN1 and NIMIN2 proteins can assemble LENRV and N1/N2BD regions of NtNPR1 in ternary complexes. It therefore seems plausible to hypothesize that NIMIN binding shapes the NPR1 C-terminus to control perception and transduction of the defense hormone SA. In fact, overexpression of *NIMIN1* severely impairs *PR* gene induction in Arabidopsis, while knock-out of *NIMIN1* enhances *PR-1* induction (Weigel *et al.*, 2005). Likewise, presence of NtNIMIN2 proteins interferes with transcription activity of the NtNPR1 C-terminus in yeast and also abolishes binding of NtTGA7 to NtNPR1 (Stos-Zweifel *et al.*, 2018). Thus, SA-induced NIMIN proteins can modulate NPR1 activity in several ways, i.e., by hampering SA perception, by concealing protein-protein interaction sites and by altering conformation in the SA-responsive NPR1 C-terminus.

## Conclusion

We propose a simple mechanism for SA-mediated activation of tobacco NPR1 (Fig. 9). In essence, our model for tobacco NPR1 is similar in some respects to a previous model for Arabidopsis NPR1. However, in contrast to the concepts formed by Wu *et al.* (2012), we find the arginine residue in the conserved LENRV motif to be essential for SA activation of the NtNPR1 C-terminus. Our model is backed by independent biochemical, genetic and physiological evidence. Furthermore, it is backed by conservation of the critical arginine residue, its immediate vicinity and a N1/N2BD generally found in NPR1 proteins. In addition, by contrast to other models, our model grants functional relevance to NPR1-interacting NIMIN proteins which, like NPR1, are present in all higher plant species. Finally, structural organization of *NPR1* genes supports our view. In Arabidopsis *NPR1*, the sequence LENR is encoded at the end of exon 2, while the core of the N1/N2BD (ELGKRFFPRCS) is encoded at the beginning of exon 4 (Fig. S14a,b). Exon 3 encompasses the whole intervening loop between LENR and the N1/N2BD core, harboring only 67 amino acids. Codons for V433 and V501 at the borders of exon 3 are generated by splicing. *NtNPR1* displays the same exon-intron structure as *AtNPR1* including codons for V432 and V499 created by excision of introns 2 and 3, respectively (Fig. S14c). This gene architecture ensures coherence of the NPR1 C-terminal region together with both conservation of LENRV and N1/N2BD motifs on the one hand and flexibility in the intervening loop on the other hand. The sequence of the intervening loop likely determines proximity of LENRV and N1/N2BD regions in the dormant protein, reach of the critical arginine into the N1/N2BD region, contact of LENRV and N1/N2BD regions in the activated protein and interplay with NIMIN proteins. It may serve as a hinge enabling differential ties between LENRV and N1/N2BD interfaces (Maier *et al.*, 2011). In this respect, it is of interest that we have found NtNPR1 proteins with mutations F505/506S in the N1/N2BD to exhibit excessive SA-mediated transcription activity in yeast, possibly suggesting altered associations in the NtNPR1 C-terminus in mutant proteins at high SA concentrations. A LENRV or LENRV-related motif with the crucial arginine residue and a N1/N2BD homologous stretch are highly conserved among NPR family members from higher plants, except BOP proteins (Maier *et al.*, 2011; Ding *et al.*, 2018). Furthermore, Arabidopsis *NPR3* and *NPR4* and tobacco *NIM1-like1* display the same exon-intron junctions as Arabidopsis and tobacco *NPR1*. Thus, NPR family members with a LENRV-like motif and a N1/N2BD-related sequence may be generally receptors for SA that perceive the SA signal in a similar manner as tobacco NPR1. The physiological outcome of structural changes in C-terminal regions, elicited by the hormone SA and modulated by NIMIN proteins, could, however, be different for diverse NPR proteins. In accord with our view, it has been shown recently that AtNPR4 and AtNPR3 function as SA-dependent transcriptional co-repressors of immune genes, acting independently from NPR1, and that SA binding and SA responsiveness are lost in mutant protein AtNPR4 R419Q (Ding *et al.*, 2018). Structural analysis of NPR1 will be needed to challenge our vision and to understand SA perception by different NPR proteins. Furthermore, structural analysis of NPR1 could provide the basis for rational design of molecular sensors for SA.

**Fig. 9.**
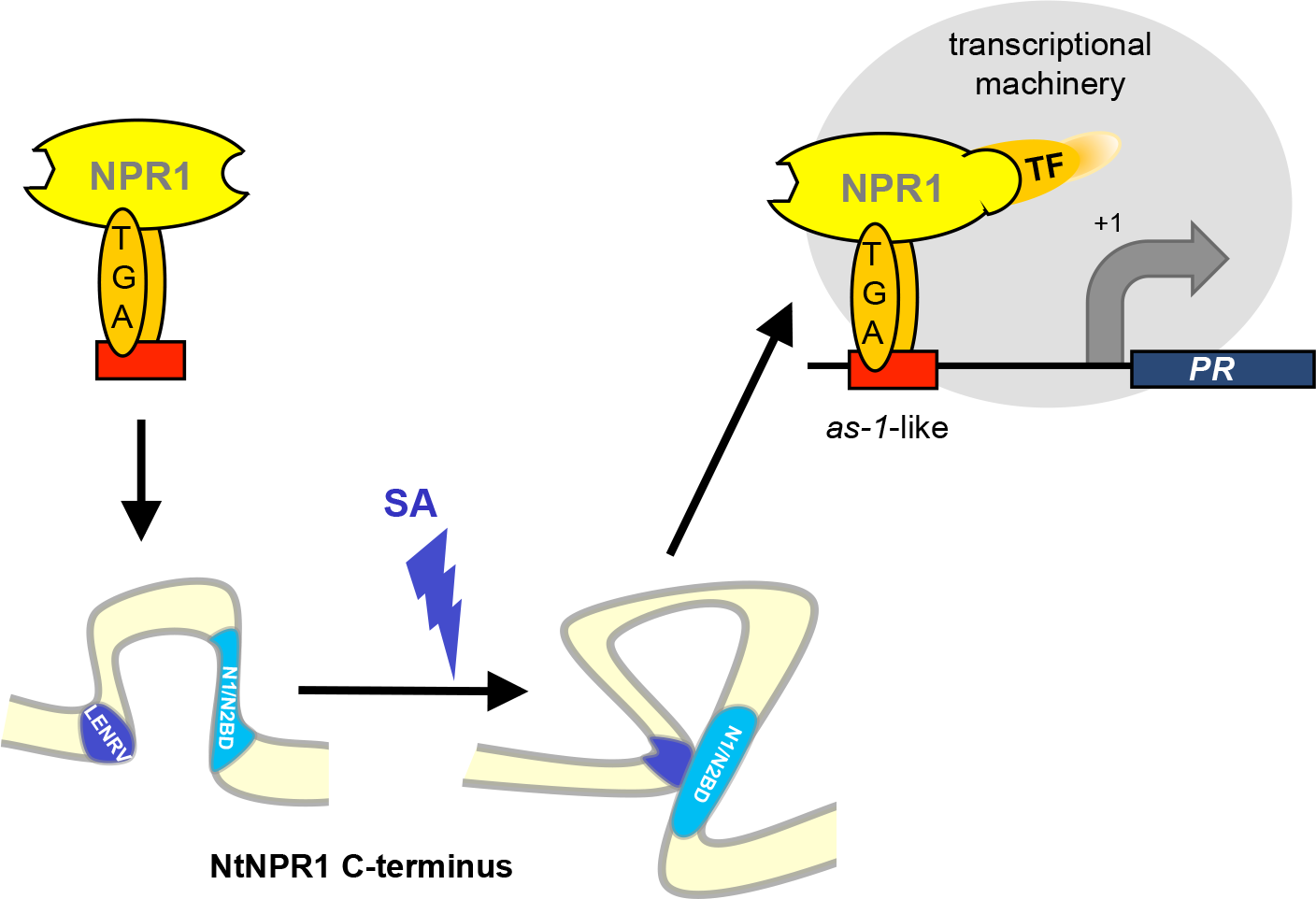
Working model for salicylic acid-driven transcription activity of the tobacco NPR1 C-terminus. The model depicts activation of the NtNPR1 C-terminus in absence of NIMIN proteins. In quiescent NPR1, LENRV and N1/N2BD regions are separated. Salicylic acid binds to the arginine side chain in the conserved LENRV motif eliciting remodeling in the NPR1 C-terminus. Contacts are built between amino acid residues of the LENRV and N1/ N2BD regions forming an SA-molded novel fold that contacts the transcription machinery to enable gene expression. Binding sites for NIMIN proteins in the N and C-terminal regions of NPR1 are indicated. N1/N2BD, binding domain for NIMIN1 and NIMIN2 proteins; TF, transcription factor.

## Acknowledgements

We thank Klaus Tietjen (Bayer CropSciences, Monheim, Germany) for the generous gift of SA analogs BTH, BTH free acid and INA; Xinnian Dong (Duke University, USA) for a tobacco *NPR1* cDNA clone; Christiane Gatz (Georg-August-Universität Göttingen, Germany) for cDNA clones encoding tobacco TGA factors 2.1, 2.2 and 10; Jörg Kudla (Westfälische Wilhelms-Universität Münster, Germany) for BiFC vectors pSPYNE-35S and pSPYCE-35S; Ashir Masroor (COMSATS University Islamabad, Pakistan) for the *NtNIMIN1-like1* clone. We also thank Christine Arnold (Universtät Hohenheim) for protein expression and pull-down experiments. This work was supported by Universität Hohenheim.

## Author contributions

DN generated gene constructs, conducted most hybrid analyses in yeast and performed transient expression of *NtNPR1* BiFC constructs in *N. benthamiana*. EK tested analogs of SA in Y2H assays and determined EC_50_ and IC_50_ values. AS performed transient expression of *NtNPR1* BiFC constructs in HEK293 cells. FM generated and analyzed mutants in the *NtNPR1* C-terminus. AJPP provided inestimable advice and fruitful discussions on the project. UMP generated gene constructs, was responsible for the coordination and supervision of the work, and wrote the article. All authors read and approved the manuscript.

## Supporting Information

Additional Supporting Information may be found online in the Supporting Information section at the end of the article.

**Fig. S1** Mutations introduced in the C-terminus of tobacco NPR1.

**Fig. S2** Controls for activity of NtNPR1 C-terminal deletions in yeast one-hybrid and two-hybrid assays.

**Fig. S3** Interaction of LENRV and N1/N2BD parts of tobacco NPR1 in yeast.

**Fig. S4** 4-Hydroxy benzoic acid (4-OH BA) does not promote physical interaction between LENRV and N1/N2BD regions of tobacco NPR1.

**Fig. S5** Visualization of LENRV and N1/N2BD interaction of tobacco NPR1 in plant cells by bimolecular fluorescence complementation (BiFC).

**Fig. S6** Association of LENRV and N1/N2BD parts of tobacco NPR1 by functional salicylic acid analogs.

**Fig. S7** Test for association of LENRV and N1/N2BD parts of tobacco NPR1 by pipecolic acid, azelaic acid and β-aminobutyric acid.

**Fig. S8** Association of LENRV and N1/N2BD parts of tobacco NPR1 by benzoic acid derivatives.

**Fig. S9** Characterization of tobacco NPR1 harboring mutation R431F.

**Fig. S10** Association of LENRV and N1/N2BD parts of tobacco NPR1 produces transcription activity in yeast.

**Fig. S11** LENRV and N1/N2BD parts of tobacco NPR1 do not interact with tobacco TGA factors.

**Fig. S12** Reconstitution of the tobacco NPR1 C-terminus from LENRV and N1/N2BD(466-525) parts by salicylic acid.

**Fig. S13** Bridging of mutant LENRV and N1/N2BD parts of tobacco NPR1 by tobacco NIMIN proteins.

**Fig. S14** Exon-intron structure in the C-terminal thirds of *NPR1* genes from Arabidopsis and tobacco.

**Table S1** Primers used for gene construction.

**Methods S1** Detailed description of methods.

