## Supplemental Information for "Salicylic acid-driven association of LENRV and NIMIN1/NIMIN2 binding domain regions in the C-terminus of tobacco NPR1 transduces SAR signal"

**Fig. S1** Mutations introduced in the C-terminus of tobacco NPR1.

**Fig. S2** Controls for activity of NtNPR1 C-terminal deletions in yeast one-hybrid and two-hybrid assays.

**Fig. S3** Interaction of LENRV and N1/N2BD parts of tobacco NPR1 in yeast.

**Fig. S4** 4-Hydroxy benzoic acid (4-OH BA) does not promote physical interaction between LENRV and N1/N2BD regions of tobacco NPR1.

**Fig. S5** Visualization of LENRV and N1/N2BD interaction of tobacco NPR1 in plant cells by bimolecular fluorescence complementation (BiFC).

**Fig. S6** Association of LENRV and N1/N2BD parts of tobacco NPR1 by functional salicylic acid analogs.

**Fig. S7** Test for association of LENRV and N1/N2BD parts of tobacco NPR1 by pipecolic acid, azelaic acid and β-aminobutyric acid.

**Fig. S8** Association of LENRV and N1/N2BD parts of tobacco NPR1 by benzoic acid derivatives.

**Fig. S9** Characterization of tobacco NPR1 harboring mutation R431F.

**Fig. S10** Association of LENRV and N1/N2BD parts of tobacco NPR1 produces transcription activity in yeast.

**Fig. S11** LENRV and N1/N2BD parts of tobacco NPR1 do not interact with tobacco TGA factors.

**Fig. S12** Reconstitution of the tobacco NPR1 C-terminus from LENRV and N1/N2BD(466-525) parts by SA.

**Fig. S13** Bridging of mutant LENRV and N1/N2BD parts of tobacco NPR1 by tobacco NIMIN proteins.

**Fig. S14** Exon-intron structure in the C-terminal thirds of *NPR1* genes from Arabidopsis and tobacco.

**Table S1** Primers used for gene construction.

**Methods S1** Detailed description of methods.

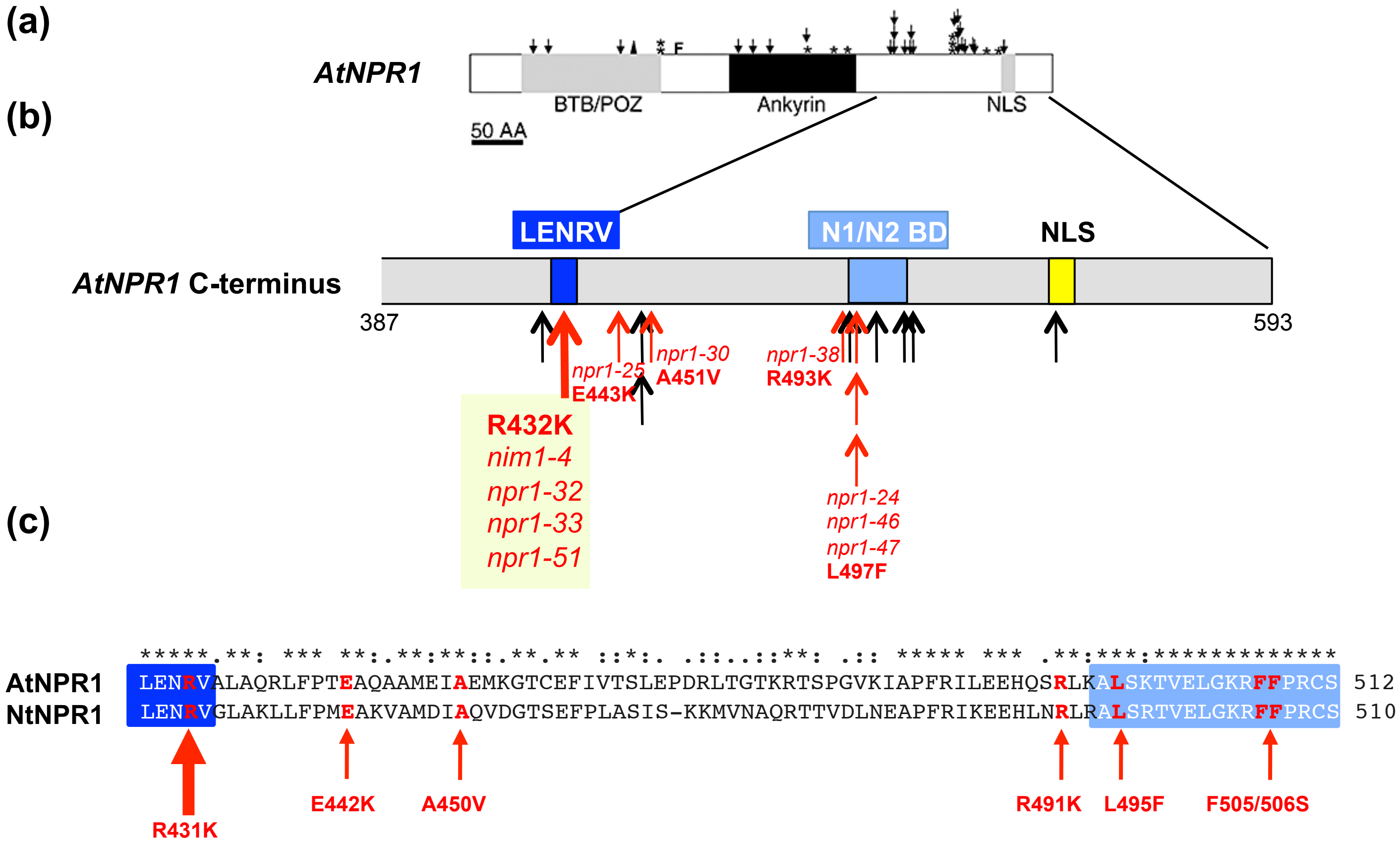


**Fig. S1** Mutations introduced in the C-terminus of tobacco NPR1. (a) Overview of Arabidopsis *npr1* alleles identified in plants insensitive to BTH. The picture was taken from Canet *et al.* (2010). Arrows denote single amino acid substitutions. Mutations are clustered in the AtNPR1 C-terminus. (b) Positions of mutations with respect to the domain structure in the Arabidopsis NPR1 C-terminus. Domains LENRV and N1/N2BD were identified by biochemical analysis and are associated with SA signaling through NPR1 proteins (Maier *et al.*, 2011). Mutations used in this study are marked in red and denoted according to Ryals *et al.* (1997; *nim1-4*) and Canet *et al.* (2010). The number of arrows at one position indicates the frequency of independent mutant alleles isolated. NLS denotes a bipartite nuclear localization signal (Kinkema *et al.*, 2000). (c) Amino acid alignment of Arabidopsis and tobacco NPR1 between the LENRV motif and the N1/N2BD. NtNPR1 mutations tested in this study are indicated. Apart from the F505/506S change, all NtNPR1 mutants are based on Arabidopsis *npr1* alleles.

**(a)**

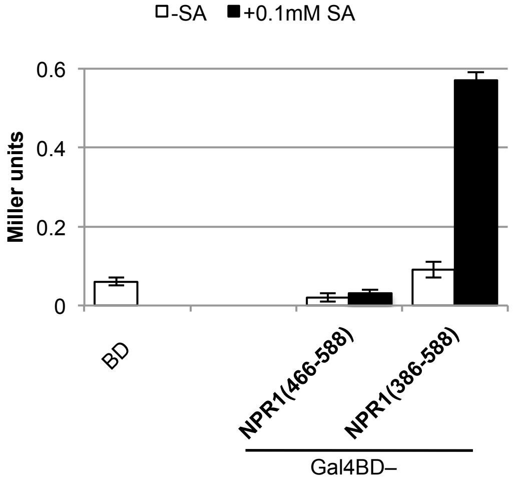


**(b)**

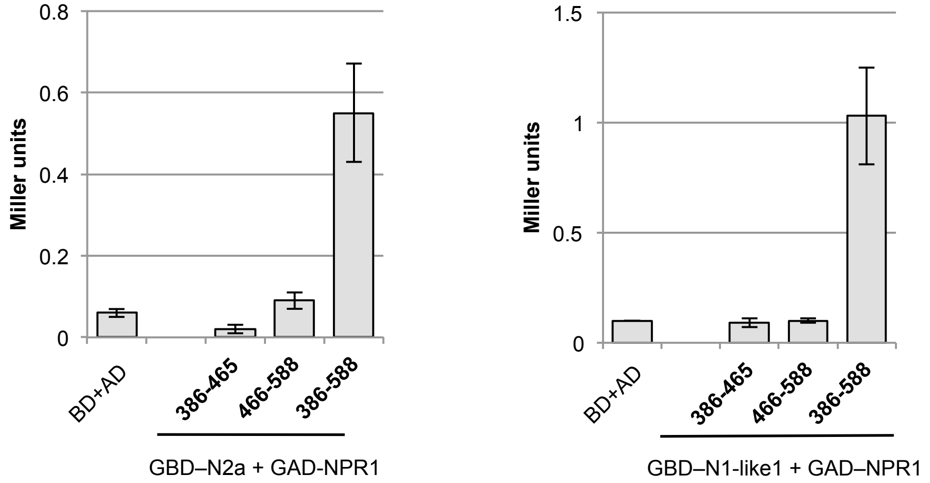


**Fig. S2** Controls for activity of NtNPR1 C-terminal deletions in yeast one-hybrid and two-hybrid assays. (a) Transcriptional activity of NtNPR1(466-588) comprising the N1/N2BD. Transactivator potential of the deleted protein was compared directly to Gal4BD–NtNPR1(386-588) in quantitative Y1H assays. Assays were conducted in absence or presence of salicylic acid. (b) Interaction of NtNIMIN2a and NtNIMIN1-like1 with NtNPR1 C-terminal deletions. Protein-protein interactions of NtNPR1(386-465) and NtNPR1(466-588) were compared directly to interaction of NtNPR1(386-588) in quantitative Y2H assays.

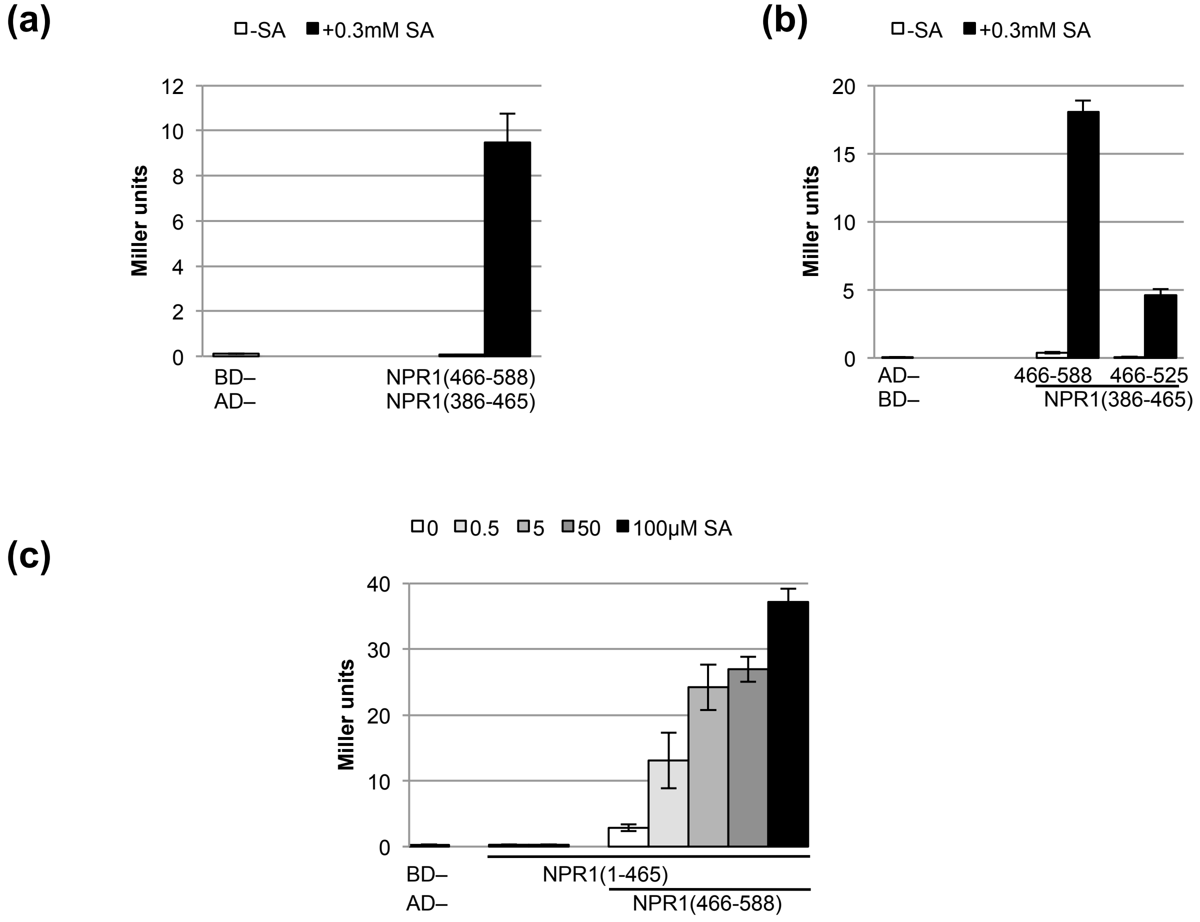


**Fig. S3** Interaction of LENRV and N1/N2BD parts of tobacco NPR1 in yeast. Protein-protein interaction in absence or presence of SA was determined in quantitative Y2H assays. (a) Interaction of GBD–NtNPR1(466-588) with GAD–NtNPR1(386-465). (b) Interaction of NtNPR1(386-465) with the NtNPR1 N1/N2BD region truncated at position 525. (c) Salicylic acid dose response curve of interaction of NtNPR1(1-465) with NtNPR1(466-588).

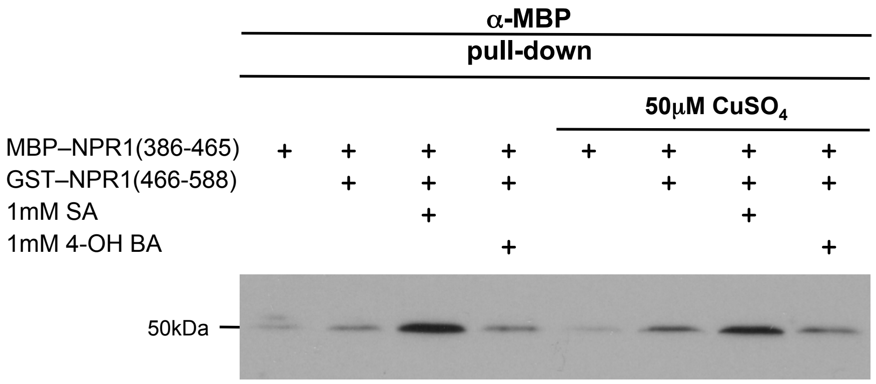


**Fig. S4** 4-Hydroxy benzoic acid (4-OH BA) does not promote physical interaction between LENRV and N1/N2BD regions of tobacco NPR1. In vitro pull-down assays of MBP–NtNPR1(386-465) with GST–NtNPR1(466-588) were performed in absence of chemicals and with SA or 4-OH BA and in presence of $Cu^{2+}$ ions.

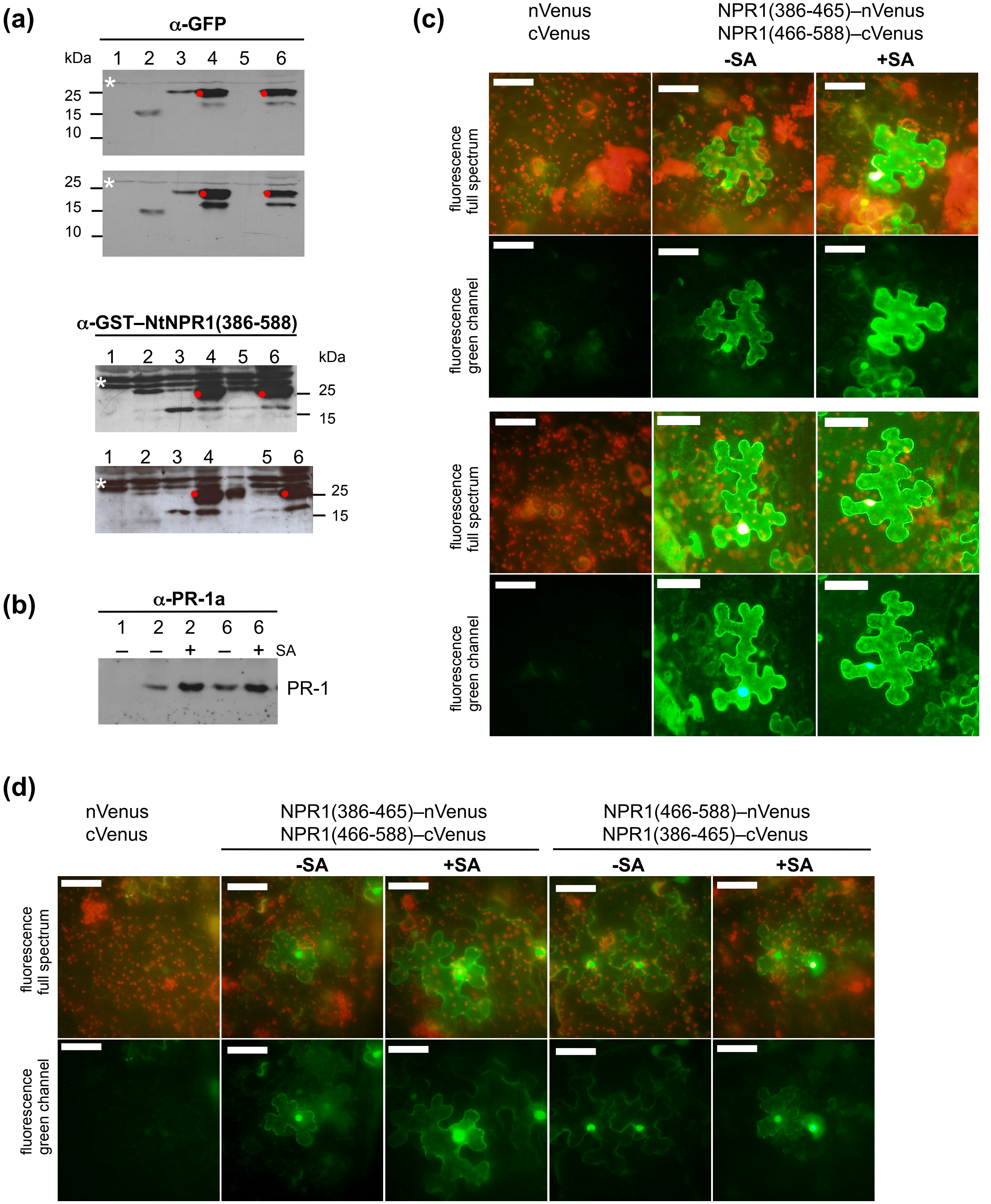
(a)
α-GFP
kDa
1 2 3 4 5 6
25
15
10
α-GST–NtNPR1(386-588)
(b)
α-PR-1a
1 2 2 6 6
– – + – + SA
PR-1
(c)
nVenus
cVenus
NPR1(386-465)–nVenus
NPR1(466-588)–cVenus
-SA
+SA
fluorescence full spectrum
fluorescence green channel
(d)
NPR1(466-588)–nVenus
NPR1(386-465)–cVenus

**Fig. S5** Visualization of LENRV and N1/N2BD interaction of tobacco NPR1 in plant cells by bimolecular fluorescence complementation (BiFC). (a) Accumulation of fusion proteins in leaves of *N. benthamiana*. Extracts were prepared from non-infiltrated leaves (lane 1), from leaves infiltrated with Agrobacteria harboring unfused *Venus* half genes (*nVenus* + *cVenus*, lane 2) or *Venus* full-length construct (lane 3) and from leaves infiltrated with Agrobacteria harboring *35S:NtNPR1(386-465)–nVenus* (*LENRV–nVenus*, lane 4), *35S:NtNPR1(466-588)–cVenus* (*N1/N2BD–cVenus*, lane 5) or both *NtNPR1* fusion genes (lane 6). Immunodetection was performed using antisera directed against GFP and against GST–NtNPR1(386-588). The NtNPR1(386-465)–nVenus fusion protein is marked by red dots. Unspecific bands detected by the antisera in any plant extract are marked by white asterisks. Results from two agroinfiltration experiments are shown. (b) Accumulation of PR-1 proteins in agroinfiltrated leaves of *N. benthamiana*. Extracts were prepared from non-infiltrated leaves (lane 1) and from leaves infiltrated with Agrobacteria as given above (lanes 2 and 6). One day prior to protein extraction, plants were sprayed with water or with 5mM SA. Immunodetection was with an antiserum directed against NtPR-1a. (c,d) BiFC analysis of LENRV and N1/N2BD interaction in salicylic acid-treated *N. benthamiana* leaf tissue and with reciprocal *Venus* half gene fusions. Leaves were infiltrated with mixtures of Agrobacterium suspensions as indicated. In panel (d), Agrobacterium strains harboring reciprocal fusion genes, *LENRV–cVenus* and *N1/N2BD–nVenus*, were used. One day prior to microscopy, plants were sprayed with water or 5mM SA. Representative results from independent agroinfiltration experiments are shown. Scale bars, 150µm.

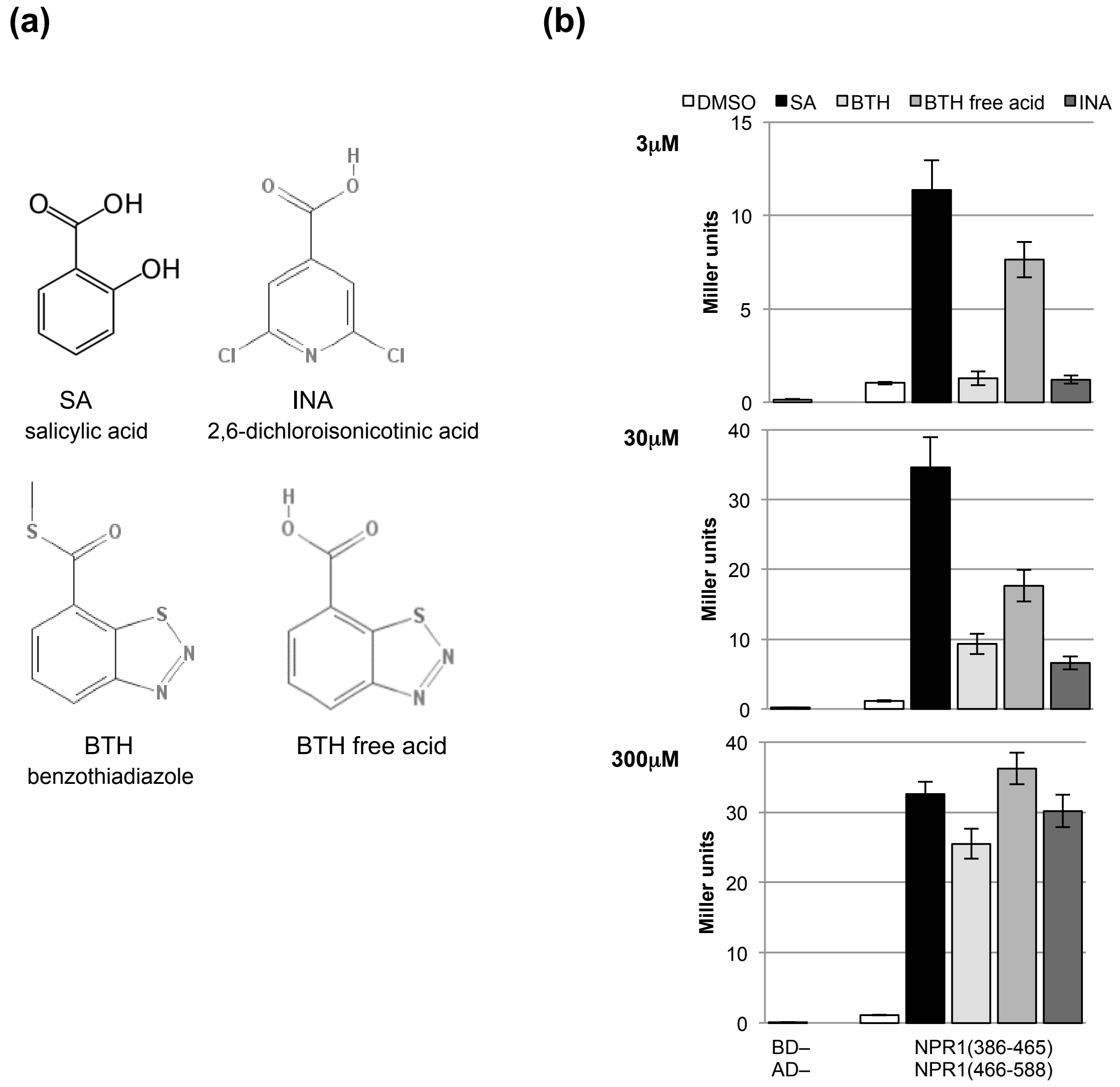


**Fig. S6** Association of LENRV and N1/N2BD parts of tobacco NPR1 by functional salicylic acid analogs. (a) Chemical structures of INA , BTH and BTH free acid (Görlach *et al.*, 1996). (b) Interaction of LENRV and N1/N2BD parts in yeast. Protein-protein interaction was determined in quantitative Y2H assays. Yeast cells were grown in medium supplemented with SA, BTH, BTH free acid or INA as indicated. Chemicals were solved in DMSO. The final DMSO concentration in yeast growth medium was 0.5%.

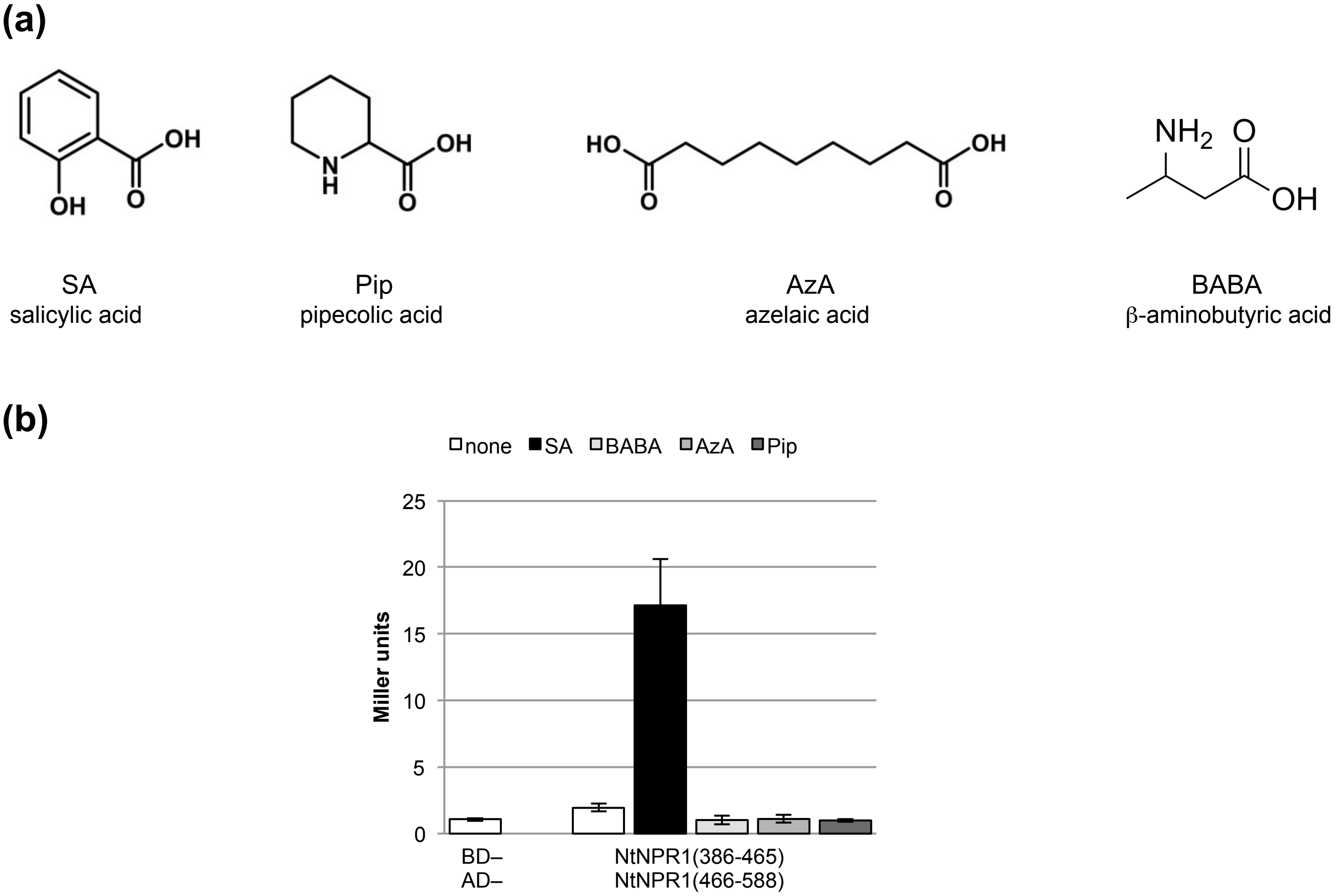


**Fig. S7** Test for association of LENRV and N1/N2BD parts of tobacco NPR1 by pipecolic acid, azelaic acid and β-aminobutyric acid. (a) Chemical structures of pipecolic acid, azelaic acid and β-amino butyric acid (Zimmerli *et al.*, 2000; Shah & Zeier, 2013). (b) Interaction of LENRV and N1/N2BD parts in yeast. Protein-protein interaction was determined in quantitative Y2H assays. Yeast cells were grown in medium supplemented with 300μM concentrations of SA or compounds as indicated. Azelaic acid was solved in DMSO. The final DMSO concentration in yeast growth medium was 0.5%.

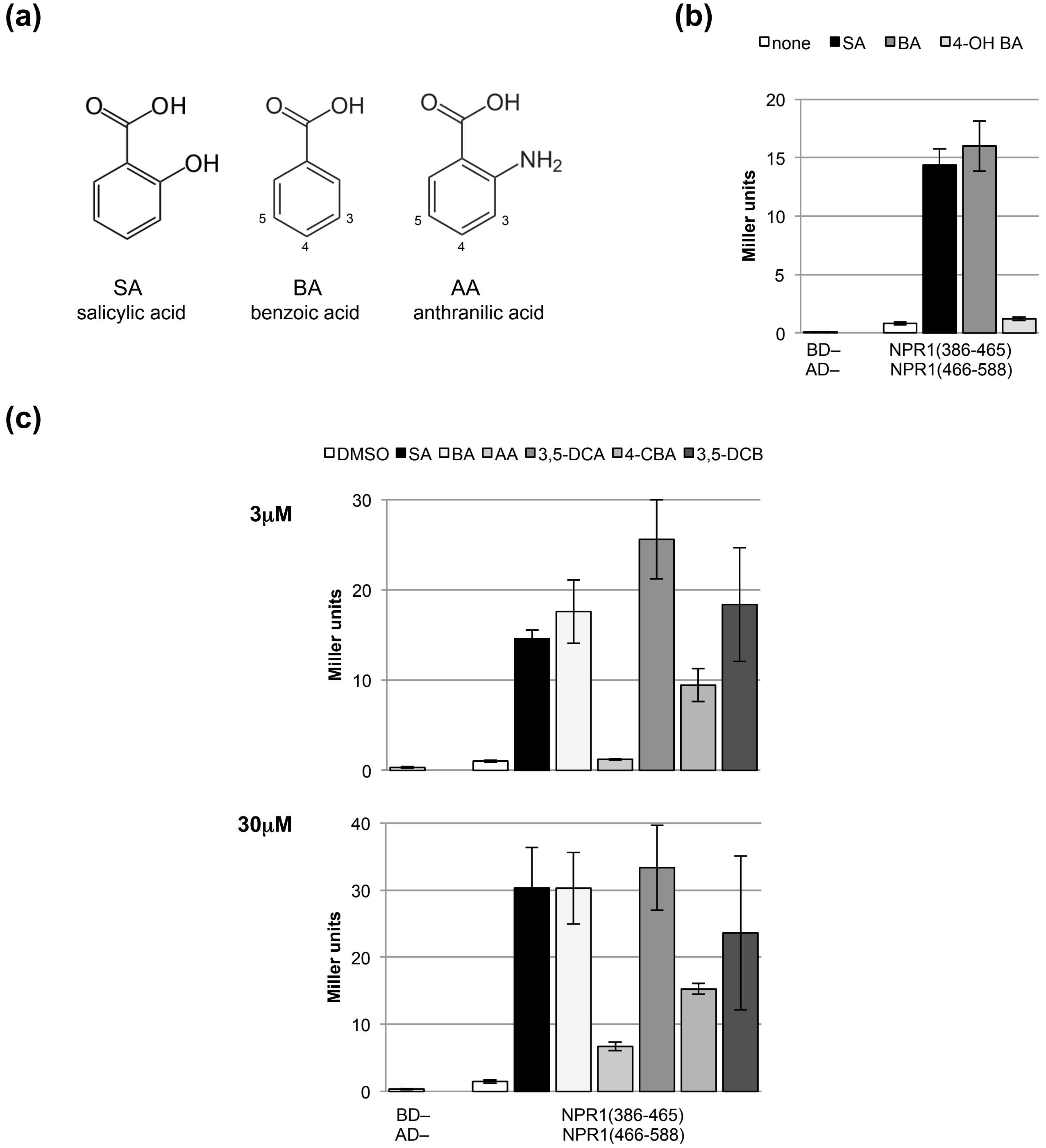


**Fig. S8** Association of LENRV and N1/N2BD parts of tobacco NPR1 by benzoic acid derivatives. (a) Chemical structures of benzoic acid and anthranilic acid. (b) Interaction of LENRV and N1/N2BD parts in presence of benzoic acid or 4-hydroxy benzoic acid (4-OH BA) in yeast. Protein-protein interaction was determined in quantitative Y2H assays. Yeast cells were grown in medium supplemented with 300µM concentrations of SA, BA or 4-OH BA. (c) Interaction of LENRV and N1/N2BD parts in presence of benzoic acid derivatives in yeast. Protein-protein interaction was determined in quantitative Y2H assays. Yeast cells were grown in medium supplemented with SA, BA, anthranilic acid (AA), 3,5-dichloroanthranilic acid (3,5-DCA), 4-chlorobenzoic acid (4-CBA) or 3,5-dichlorobenzoic acid (3,5-DCB) as indicated. Chemicals were solved in DMSO. The final DMSO concentration in yeast growth medium was 0.5%.

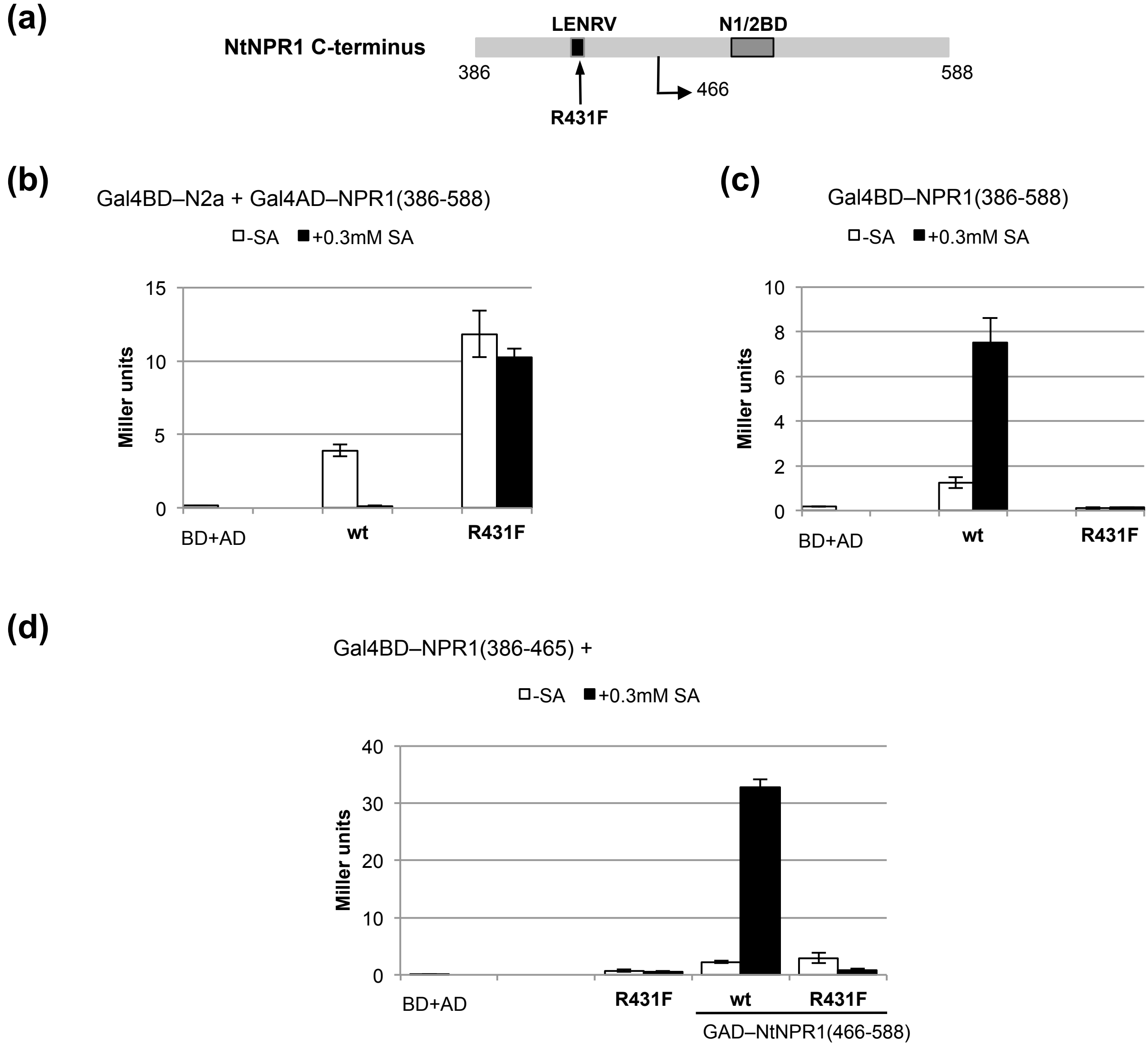


**Fig. S9** Characterization of tobacco NPR1 harboring mutation R431F. Quantitative Y1H and Y2H assays were performed with cells grown in medium without or with 300µM SA. (a) Position of mutation R431F in the tobacco NPR1 C-terminus. (b) Interaction of NtNPR1(386-588)R431F with tobacco NIMIN2a. (c) Transcription activity of NtNPR1(386-588)R431F. (d) Interaction of LENRV R431F and N1/N2BD parts.

**(a)**

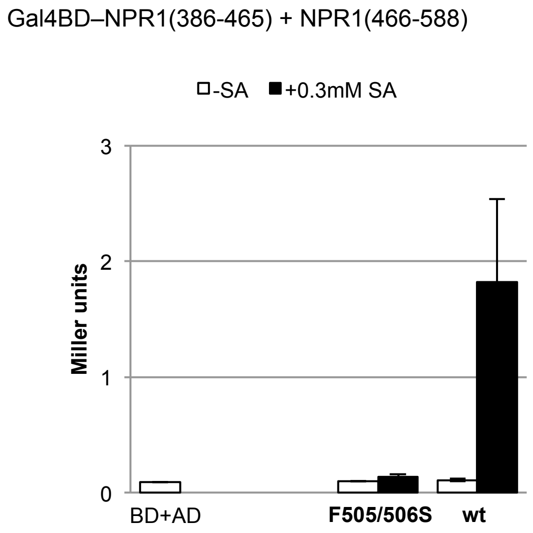


**(b)**

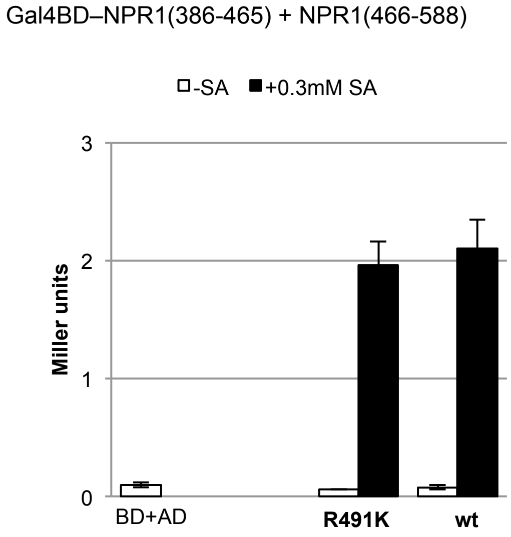


**Fig. S10** Association of LENRV and N1/N2BD parts of tobacco NPR1 produces transcription activity in yeast. Quantitative Y2H assays were performed with cells grown in medium without or with 300μM SA. (a) Mutant NtNPR1(466-588)F505/506S does not reconstitute a transcriptionally active C-terminus. (b) Mutant NtNPR1(466-588)R491K reconstitutes a transcriptionally active C-terminus.

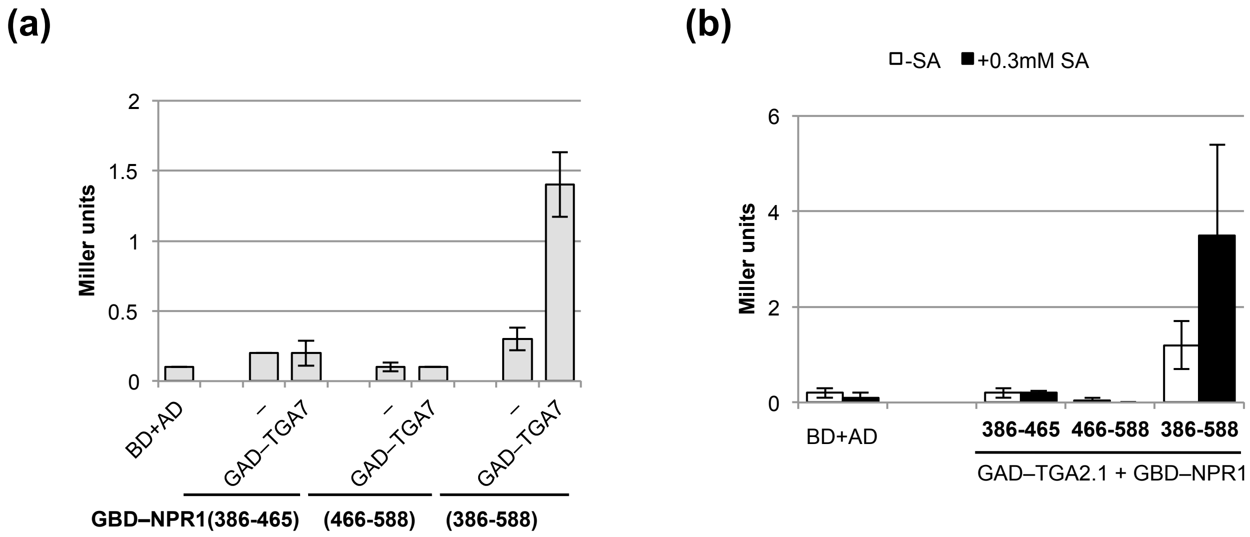


**Fig. S11** LENRV and N1/N2BD parts of tobacco NPR1 do not interact with tobacco TGA factors. Quantitative Y2H interaction of NtNPR1(386-465), NtNPR1(466-588) and NtNPR1(386-588) with tobacco TGA7 (a) and TGA2.1 (b).

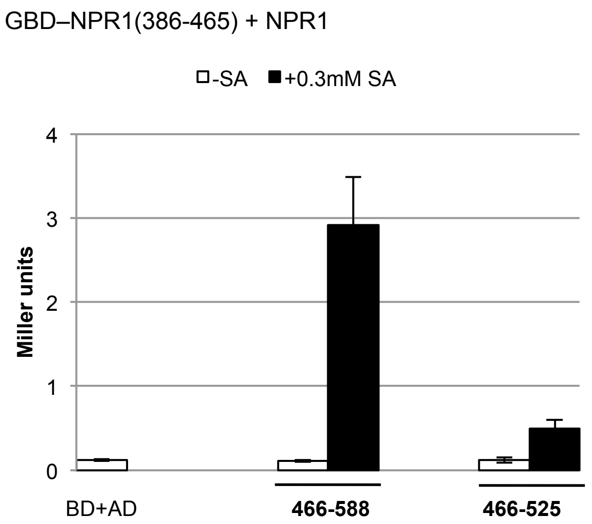


**Fig. S12** Reconstitution of the tobacco NPR1 C-terminus from LENRV and N1/N2BD (466-525) parts by salicylic acid. *N1/N2BD* parts were expressed from the *MET25* promoter. Quantitative protein-protein interaction assays were performed with cells grown in medium lacking methionine without or with SA. Transcription activity of the reconstituted C-terminus truncated at amino acid 525 was compared to the full NtNPR1 C-terminus reconstituted from LENRV and N1/N2BD(466-588) parts.

**(a)**

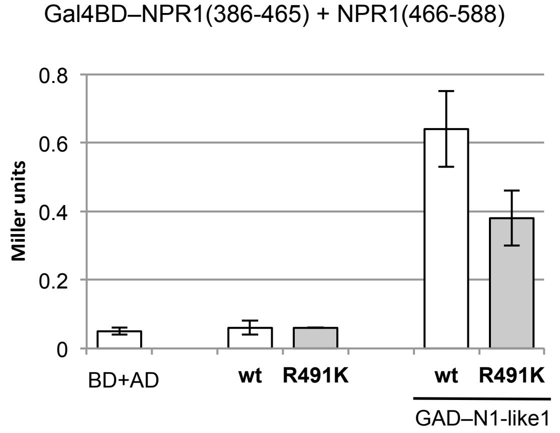


**(b)**

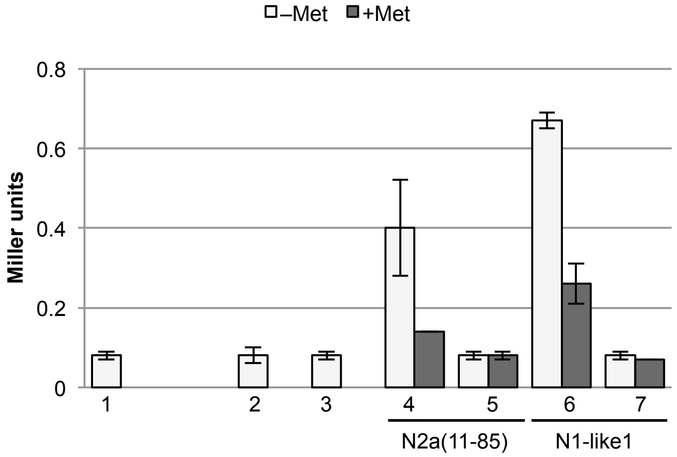


| | GBD fusion | GAD fusion | Conditional (±Met) |
|---|---|---|---|
| **1:** | – | – | – |
| **2:** | NPR1(386-465) | – | NPR1(466-588) |
| **3**: | NPR1(386-465) R431K | – | NPR1(466-588) |
| **4**: | NPR1(386-465) | N2a (11-85) | NPR1(466-588) |
| **5**: | NPR1(386-465) R431K | N2a (11-85) | NPR1(466-588) |
| **6**: | NPR1(386-465) | N1-like1 | NPR1(466-588) |
| **7**: | NPR1(386-465) R431K | N1-like1 | NPR1(466-588) |

**Fig. S13** Bridging of mutant LENRV and N1/N2BD parts of tobacco NPR1 by tobacco NIMIN proteins. (a) Tobacco NIMIN1-like1 can bridge LENRV and N1/N2BD R491K parts of tobacco NPR1. Quantitative Y3H interaction assays were performed with cells grown in absence of methionine. (b) Tobacco NIMIN proteins cannot bridge LENRV R431K and N1/N2BD parts of tobacco NPR1. Quantitative yeast hybrid interaction assays were performed with cells grown in absence and presence of methionine.

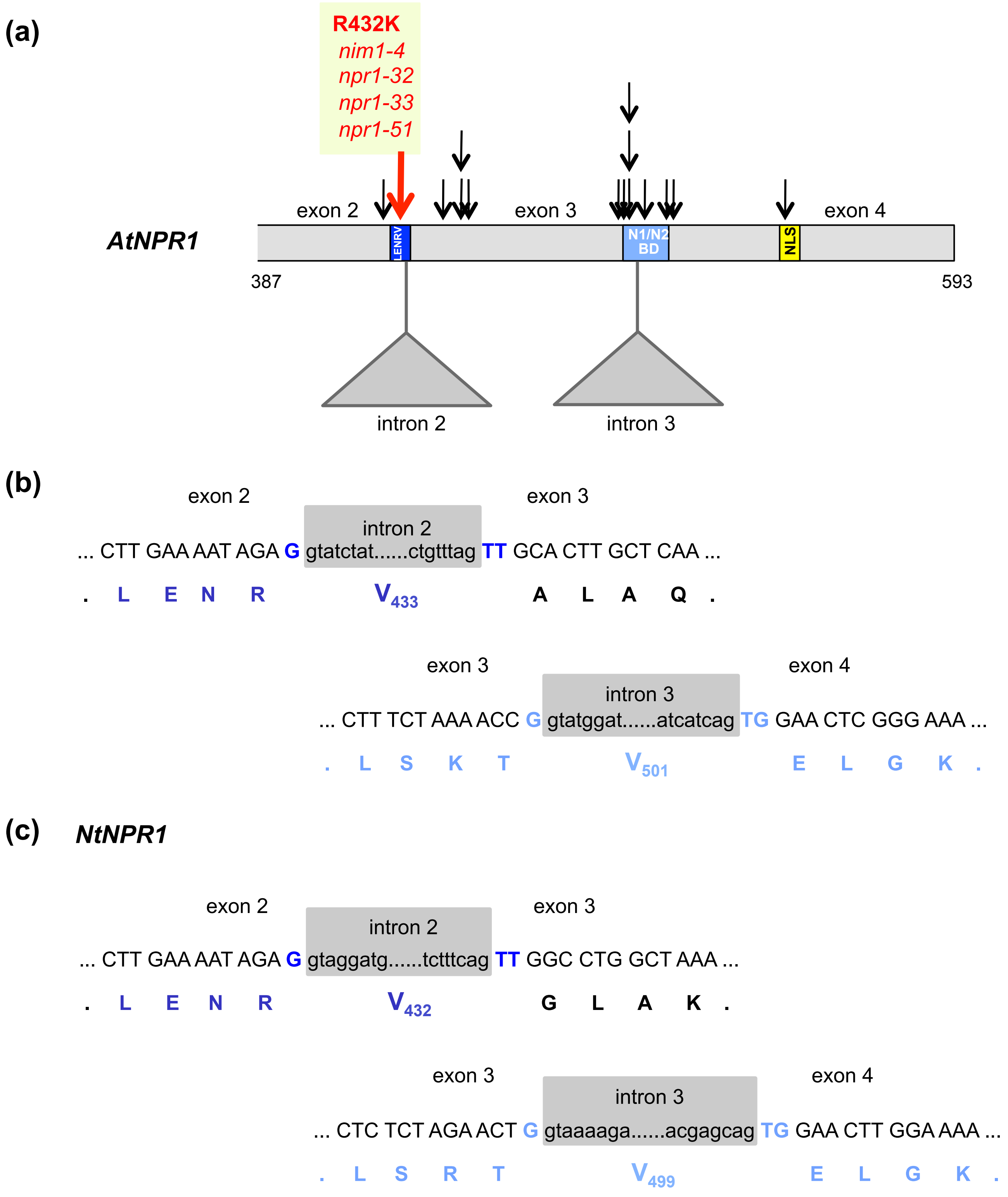


**Fig. S14** Exon-intron structure in the C-terminal thirds of *NPR1* genes from Arabidopsis and tobacco. (a) Exon-intron structure in the C-terminal third of Arabidopsis *NPR1*. Positions of introns 2 and 3 are depicted with respect to the LENRV motif and the N1/N2BD. Single amino acid substitutions identified in BTH-insensitive *npr1* alleles in planta are marked by arrows (Canet *et al.*, 2010). Exons 3 and 4 are drawn approximately to scale. (b) Splice junctions of introns 2 and 3 of the Arabidopsis *NPR1* gene. The codon for V433 in the conserved LENRV motif is encoded by exons 2 and 3, and the codon for V501 in the N1/N2BD is encoded by exons 3 and 4. (c) Splice junctions of introns 2 and 3 of the tobacco *NPR1* gene. The codon for V432 in the conserved LENRV motif is encoded by exons 2 and 3, and the codon for V499 in the N1/N2BD is encoded by exons 3 and 4. N1/N2BD, binding domain for NIMIN1 and NIMIN2 proteins; NLS, nuclear localization signal.

**Table S1.** Primers used for gene construction.

| Construct | Primer name | Sequence 5´ to 3´ |
|---|---|---|
| *NtNPR1(386-588)* | tNPR1-10 | AAGGATCCGTTCTGCTTCGAATGATCGG |
| | tNPR1-2 | TTGTCGACCTATTTCCTAAAAGGGAGC |
| *E442K* | tNPR1-23 | CTCCTTTTTCCAATGAAAGCTAAAGTTGCAATGG |
| | tNPR1-24 | CCATTGCAACTTTAGCTTTCATTGGAAAAAGGAG |
| *A450V* | tNPR1-16 | GCTAAAGTTGCAATGGACATTGTTCAAGTTGATGGCACTTCTGAG |
| | tNPR1-17 | CTCAGAAGTGCCATCAACTTGAACAATGTCCATTGCAACTTTAGC |
| *L495F* | tNPR1-18 | GAATCGGCTTAGAGCATTCTCTAGAACTGTAGAACTTGG |
| | tNPR1-19 | CCAAGTTCTACAGTTCTAGAGAATGCTCTAAGCCGATTC |
| *R491K* | tNPR1-29 | TTCTAGAGAGTGCTCTAAGCTTATTCAAGTGCTCCTC |
| *LENRV(386-465)* | tNPR1-22 | AAGTCGACTTTGCCGATGCTAGCCAGTGGG |
| *N1/N2BD(466-588)* | tNPR1-21 | TTGGATCCAAAAGATGGCTAATGCACAGAGG |
| *N1/N2BD(466-525)* | pGADback | GCATGCCGGTAGAGGTGT |
| *Venus* | V1a | GCCCGGGGGATCCACCATGGTGAGCAAGGGCGAG |
| | V1b | GGAGCTCTTACTTGTACAGCTCGTCCATG |
| *nVenus* | Venus-1 | CCCGGGATGGTGAGCAAGGGCGAGGAGC |
| | Venus-2 | GAGCTCTTAGGTGATATAGACGTTGTGG |
| *cVenus* | Venus-3 | GCCCGGGATGGCCGACAAGCAGAAGAACGG |
| | Venus-4 | GGGAGCTCTTACTTGTACAGCTCG |
| *vNtNPR1(386-465)* | ctNPR1-1 | GGATCCAAATGTCTGCTTCGAATGATCGG |
| | ctNPR1-2 | GGTCGACTTACTCGAGTTTGCCGATGCTAGCCAGTGGG |
| *vNtPR1(466-588)* | ctNPR1-3 | GGATCCAAATGAAGATGGCTAATGCACAGAGG |
| | ctNPR1-4 | GGTCGACCTACTCGAGTTTCCTAAAAGGGAGCTTATTGGG |

**Methods S1** Detailed description of methods.

### DNA constructs

For protein interaction assays in yeast, cDNA sequences encoding truncated or full-length proteins were fused in-frame to the sequence for GBD in pGBT9 and the sequence for GAD in pGAD424. For expression of a third gene in Y3H interaction analyses, gene sequences were excised from pGBT9 and pGAD424 constructs as *Bam*HI or *Bam*HI/*Bgl*II fragments and inserted in the *Bgl*II restriction enzyme site of a modified pBridge vector, designated pBD, under control of the *MET25* promoter (Weigel *et al.*, 2001). Clones containing full-length or partial *NIMIN* cDNAs have been described previously (Zwicker *et al.*, 2007). All clones generated by PCR amplification were verified by DNA sequence analysis. Primers used for cloning are listed in Table S1.

Generation of *NtNPR1* full-length and partial (aa 386 to 588 and aa 386 to 525) gene constructs and cloning of mutants *NtNPR1(386-588)R431K*, *NtNPR1 F505/506S* and *NtNPR1(386-588)F505/506S* have been described earlier (Maier *et al.*, 2011; Stos-Zweifel *et al.*, 2018). Additional mutants used in this work were generated by overlap extension PCR in the *NtNPR1* cDNA encoding amino acids 386 to 588 (Ho *et al.*, 1989). tNPR1-10 and tNPR1-2 were used as forward and back primers, respectively, in combination with various mutagenesis primers (Table S1). PCR products were cloned as *Bam*HI/*Sal*I fragments into vectors pGBT9 and pGAD424. Mutant R491K was generated by PCR with mutagenesis primer tNPR1-29 harboring an *Xba*I restriction endonuclease site. The 0.3 kb N-terminal *Bam*HI/*Xba*I fragment in pGAD424/NtNPR1(386-588) was replaced with the fragment carrying the R491K mutation. Partial *NtNPR1* cDNA sequences encoding LENRV (aa 386 to 465) and N1/N2BD (aa 466 to 588) regions were generated by PCR from wild-type and mutant pGBT9/NtNPR1(386-588) constructs using primer pairs tNPR1-10/tNPR1-22 and tNPR1-21/tNPR1-2, respectively. The sequence encoding the deleted N1/N2BD region from amino acids 466 to 525 was amplified from pGBT9/NtNPR1(386-525) with primers tNPR1-21 and pGADback. The fragment was cloned into *Bam*HI cut pGAD424 and into *Bgl*II cut pBD. Construction of the full-length *LENRV* sequence from amino acids 1 to 465 was described by Stos-Zweifel *et al.* (2018). For expression of fusion genes in *E. coli*, the *LENRV* sequence was excised with *Eco*RI and *Sal*I from pGBT9/NtNPR1(386-465) and its mutant version R431K. The fragments were inserted in *Eco*RI/*Sal*I cleaved expression vector pMAL-c2 to produce a translational fusion at the C-terminus of maltose binding protein (MBP). Similarly, the *N1/N2BD* sequence was excised with *Eco*RI and *Sal*I from pGAD424/NtNPR1(466-588), and the fragment was inserted in *Eco*RI/*Sal*I cleaved expression vector pGEX-5X to produce a translational fusion at the C-terminus of glutathione S-transferase (GST).

BiFC constructs to investigate LENRV–N1/N2BD interaction in planta were assembled in modified binary vectors pSPYNE-35S and pSPYCE-35S (Walter *et al.*, 2004). *Yellow fluorescent protein* (*YFP*) half genes including epitope tags were removed from the vectors through *Sma*I/*Sac*I digestion and replaced by *Venus* half genes (Nagai *et al.*, 2002) amplified with primer pairs Venus-1/Venus-2 (aa 1-154, nVenus) and Venus-3/Venus-4 (aa 155-239, cVenus). Fragment *nVenus* was ligated to pSPYNE-35S, and fragment *cVenus* was transferred to pSPYCE-35S yielding cloning vectors pSPYNE2 and pSPYCE2. *LENRV* and *N1/N2BD* parts of *NtNPR1* were amplified by PCR using primer pairs ctNPR1-1/ctNPR1-2 and ctNPR1-3/ctNPR1-4, respectively. PCR products contained ATG translation start codons and stop codons 3´ to an *Xho*I restriction endonuclease site. Fragments with *Bam*HI/*Xho*I ends were cloned into *Bam*HI/*Xho*I cleaved pSPYNE2 and pSPYCE2 to produce translational fusions at the 3´-ends of *NtNPR1* parts with *Venus* half genes. As a positive control, pBin19/35S:Venus was used. The *Venus* sequence was amplified with primers V1a and V1b. The resulting fragment was cut with *Bam*HI and *Sac*I and transferred to pBin19/35S:GUS (Jefferson *et al.*, 1987) replacing the *GUS* reporter. *Venus* and *Venus* half genes were expressed from the *CaMV 35S* promoter. The basis for expression of *NtNPR1-Venus* half genes in animal cells was pEYFP-Mem (CLONTECH Laboratories). The *EYFP* gene including the membrane attachment signal was removed from the vector through digestion with *Age*I and *Not*I. Overhanging ends were filled in with Klenow DNA polymerase, and the vector was religated to yield pΔEYFP-Mem. *NtNPR1-Venus* half genes from pSPYNE2 and pSPYCE2 vectors were transferred as *Bam*HI/*Sac*I fragments to pΔEYFP-Mem cleaved with *Bgl*II and *Sac*I. As negative controls, unfused *Venus* half genes isolated as *Xho*I/*Sac*I fragments from pSPYNE2 and pSPYCE2 vectors were cloned in *Xho*I/*Sac*I cleaved pΔEYFP-Mem. Expression of *Venus* half genes is driven by the immediate early promoter of CMV.

**Accession numbers**

Sequence data from this article can be found in the EMBL/GenBank databases under the following accession numbers:

AF480488 (*NtNPR1*), NW_015861707 (*NtNPR1* gene), AF057379 (*NtNIMIN2a*), EF015598 (*NtNIMIN2c*), BP531936 (*NtNIMIN1-like1*), KY392756 (*NtTGA7*), X16449 (*NtTGA1a*), U90214 (*NtTGA2.1*), AF031487 (*NtTGA2.2*), AY998018 (*NtTGA10*).


**Canet JV, Dobón A, Roig A, Tornero P. 2010.** Structure-function analysis of npr1 alleles in Arabidopsis reveals a role for its paralogs in the perception of salicylic acid. *Plant, Cell and Environment* **33**: 1911-1922.

**Ho SN, Hunt HD, Horton RM, Pullen JK, Pease LR. 1989.** Site-directed mutagenesis by overlap extension using the polymerase chain reaction. *Gene* **77**: 51-59.

**Görlach J, Volrath S, Knauf-Beiter G, Hengy G, Beckhove U, Kogel KH, Oostendorp M, Staub T, Ward E, Kessmann H *et al.* 1996.** Benzothiadiazole, a novel class of inducers of systemic acquired resistance, activates gene expression and disease resistance in wheat. *Plant Cell* **8**: 629-643.

**Jefferson RA, Kavanagh TA, Bevan MW. 1987.** GUS fusions: beta-glucuronidase as a sensitive and versatile gene fusion marker in higher plants. *EMBO Journal* **6**: 3901-3907.

**Kinkema M, Fan W, Dong X. 2000.** Nuclear localization of NPR1 is required for activation of PR gene expression. *Plant Cell* **12**: 2339-2350.

**Maier F, Zwicker S, Hückelhoven A, Meissner M, Funk J, Pfitzner AJP, Pfitzner UM. 2011.** NONEXPRESSOR OF PATHOGENESIS-RELATED PROTEINS1 (NPR1) and some NPR1-related proteins are sensitive to salicylic acid. *Molecular Plant Pathology* **12**: 73-91.

**Nagai T, Ibata K, Park ES, Kubota M, Mikoshiba K, Miyawaki A. 2002.** A variant of yellow fluorescent protein with fast and efficient maturation for cell-biological applications. *Nature Biotechnology* **20**: 87-90.

**Ryals J, Weymann K, Lawton K, Friedrich L, Ellis D, Steiner HY, Johnson J, Delaney TP, Jesse T, Vos P *et al.* 1997.** The Arabidopsis NIM1 protein shows homology to the mammalian transcription factor inhibitor I kappa B. *Plant Cell* **9**: 425-439.

**Shah J, Zeier J. 2013.** Long-distance communication and signal amplification in systemic acquired resistance. *Frontiers in Plant Science* **4**: 30.

**Stos-Zweifel V, Neeley D, Konopka E, Meissner M, Hermann M, Maier F, Häfner V, Pfitzner AJP, Pfitzner UM. 2018.** Tobacco TGA7 mediates gene expression dependent and independent of salicylic acid. *Bioarchives.* doi: 10.1101/341834

**Walter M, Chaban C, Schütze K, Batistic O, Weckermann K, Näke C, Blazevic D, Grefen C, Schumacher K, Oecking C *et al.* 2004.** Visualization of protein interactions in living plant cells using bimolecular fluorescence complementation. *The Plant Journal* **40**: 428-438.

**Weigel RR, Bäuscher C, Pfitzner AJP, Pfitzner UM. 2001.** NIMIN-1, NIMIN-2 and NIMIN-3, members of a novel family of proteins from Arabidopsis that interact with NPR1/NIM1, a key regulator of systemic acquired resistance in plants. *Plant Molecular Biology* **46**: 143-160.

**Zimmerli L, Jakab G, Métraux J-P, Mauch-Mani B. 2000.** Potentiation of pathogen-specific defense mechanisms in Arabidopsis by β-aminobutyric acid. *Proceedings of the National Academy of Sciences, USA* **97**: 12920-12925.

**Zwicker S, Mast S, Stos V, Pfitzner AJP, Pfitzner UM. 2007.** Tobacco NIMIN2 proteins control PR gene induction through transient repression early in systemic acquired resistance. *Molecular Plant Pathology* **8**: 385-400.
